# Transposon-mediated genic rearrangements underlie variation in small RNA pathways

**DOI:** 10.1101/2024.01.15.575659

**Authors:** Gaotian Zhang, Marie-Anne Félix, Erik C. Andersen

## Abstract

Transposable elements (TEs) are parasitic DNA sequences that insert into the host genome and can cause alterations in host gene structure and expression. Host organisms cope with the often detrimental consequences caused by recent transposition and develop mechanisms that repress TE activities. In the nematode *Caenorhabditis elegans*, a small interfering RNA (siRNA) pathway dependent on the helicase ERI-6/7 primarily silences long terminal repeat retrotransposons and recent genes of likely viral origin. By studying gene expression variation among wild *C. elegans* strains, we discovered that structural variants and transposon remnants at the *eri-6*/*7* locus alter its expression in *cis* and underlie a *trans*-acting expression quantitative trait locus affecting non-conserved genes and pseudogenes. Multiple insertions of the *Polinton* DNA transposon (also known as *Mavericks*) reshuffled the *eri-6/7* locus in different configurations, separating the *eri-6* and *eri-7* exons and causing the inversion of *eri-6* as seen in the reference N2 genome. In the inverted configuration, gene function was previously shown to be repaired by unusual *trans*-splicing mediated by direct repeats flanking the inversion. We show that these direct repeats originated from terminal inverted repeats specific to *C. elegans Polintons*. This *trans*-splicing event occurs infrequently compared to *cis*-splicing to novel downstream exons, thus affecting the production of ERI-6/7. Diverse *Polinton*-induced structural variations display regulatory effects within the locus and on targets of ERI-6/7-dependent siRNA pathways. Our findings highlight the role of host-transposon interactions in driving rapid host genome diversification among natural populations and shed light on evolutionary novelty in genes and splicing mechanisms.

## Main

Transposable elements (TEs) are ubiquitous mobile DNA sequences. With their parasite-like nature and the invasive mechanisms of transposition, these selfish genetic elements propagate in host genomes and cause diverse mutations, ranging from point mutations to genome rearrangements and expansions^1–3^. They can even transfer horizontally across individuals and species, leading to movement of genetic material between widely diverged taxa^4,5^. To the hosts, recent TE insertions are mostly deleterious. Various pathways have evolved in hosts to repress expression and transposition of TEs^6–9^. By contrast, hosts can also benefit from TEs, because TE sequences can serve as building blocks for the emergence of protein-coding genes, non-coding RNAs, centromeres, and *cis*-regulatory elements^10–12^.

Small RNAs are widely used to repress expression of TEs and other genes^6,7,9^. In the nematode *Caenorhabditis elegans*, the helicase ERI-6/7-dependent small interfering RNAs (siRNAs) primarily target long terminal repeat (LTR) retrotransposons and pairs or groups of non-conserved genes and pseudogenes that show extensive homology and have likely viral origins^9,13^. The closest known species of *C. elegans*, *Caenorhabditis inopinata*, lost the *eri-6/7* related small RNA pathway, which was suggested to have caused the expansion of transposons in its genome compared to *C. elegans* and another related species, *Caenorhabditis briggsae*^9,14^. In *C. elegans*, ERI-6/7 is required for the biogenesis of the Argonaute ERGO-1-associated endogenous siRNAs (Fig. 1a)^13^. Likely because endogenous and exogenous siRNA pathways share and compete for downstream resources^15^, mutants of *eri-6/7* display enhanced RNA interference (RNAi) responses to exogenous dsRNAs^16^. Competition also exists among different endogenous siRNA pathways. Within the *eri-6*/*7* locus, three other local open reading frames (*eri-6[e]*, *eri-6[f]*, and *sosi-1*) act independently of one another in a feedback loop to modulate the expression of ERI-6/7 and maintain a balance between different endogenous siRNAs (Fig. 1a, b)^17^.

**Fig. 1:**
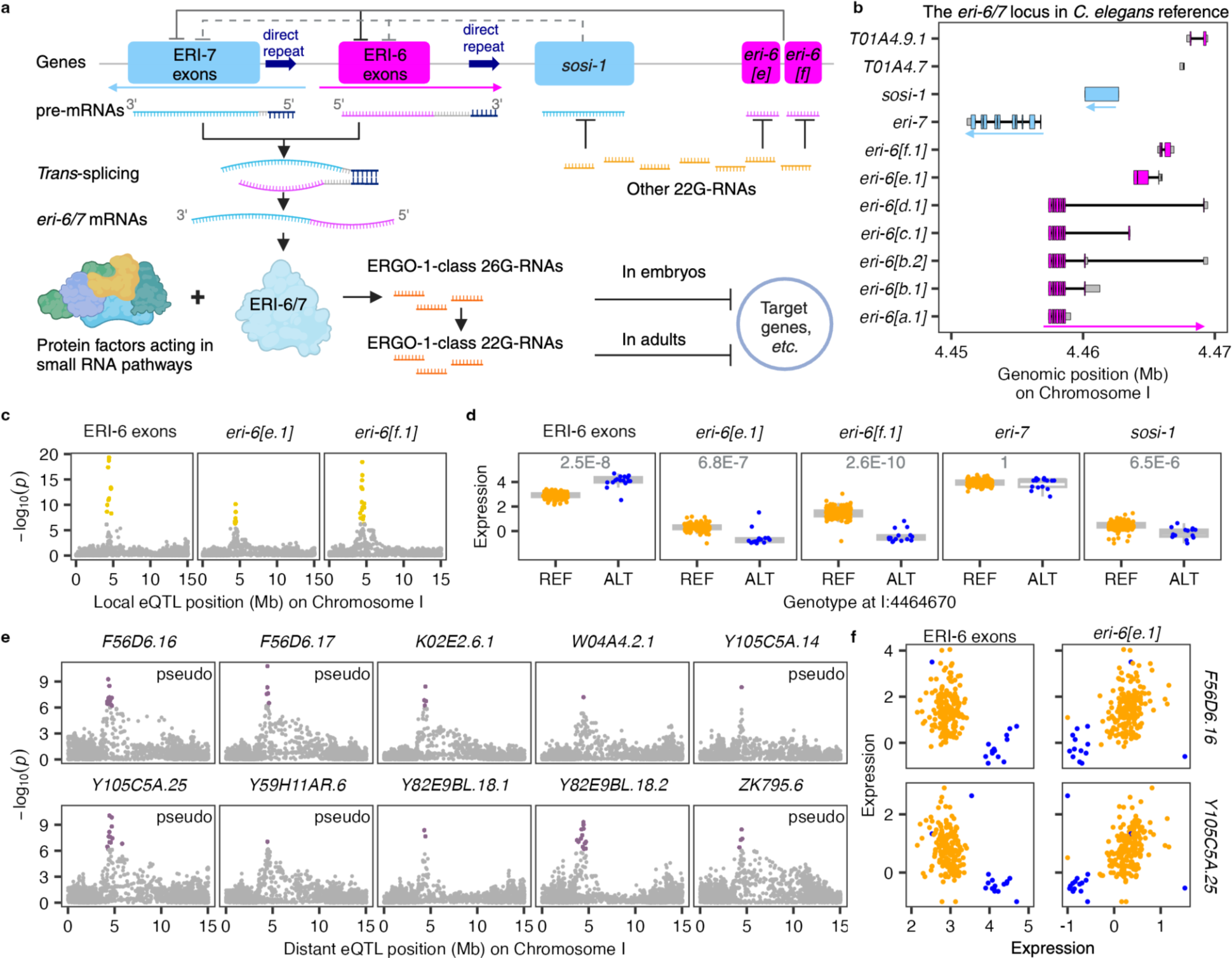
Expression variation in *eri-6* potentially mediates a *trans*-acting eQTL hotspot. **a**, Graphic illustration of the ERI-6/7-dependent siRNA pathways and the feedback loop. Dark blue arrows indicate direct repeats. Pink and blue rectangles indicate exons on the plus and minus strand, respectively (The same color scheme is used in the following figures). Created using BioRender. **b**, Structures of genes and isoforms at the *eri-6/7* locus in the reference genome (WS283)^22^. **c**, **e**, Manhattan plots indicating the GWAS mapping results of transcript expression traits on chromosome I for ERI-6 exons, *eri-6[e]*, and *eri-6[f]* (**c**) and ten transcripts across the genome (**e**). Each point represents a SNV that is plotted with its genomic position (x-axis) against its -log10(*p*) value (y-axis) in mappings. SNVs that pass the 5% FDR threshold are colored gold and purple for local and distant eQTL, respectively. Transcripts of pseudogenes are indicated. **d**, Tukey box plots showing expression (-*log*_2_(normalized TPM+0.5)) variation of five transcripts at the *eri-6/7* locus between strains with different alleles at the top candidate SNV (I: 4,464,670). Statistical significance of each comparison is shown above and was calculated using the two-sided Wilcoxon test and was corrected for multiple comparisons using the Bonferroni method. **f**, Correlations of expression variation of two transcripts to expression variation of ERI-6 exons and *eri-6[e]*. Each point (**d, f**) represents a strain and is colored orange and blue for strains with the reference (REF) or the alternative (ALT) allele at the SNV, respectively.

In addition to the vital role of ERI-6/7 in RNAi pathways, its discovery^16^ revealed a highly unusual expression mechanism. Fischer and Ruvkun showed that *eri-6* and *eri-7*, two adjacent genes oriented in opposing genomic directions in the *C. elegans* reference strain N2, employ a *trans*-splicing mechanism to generate fused *eri-6/7* mRNAs encoding the helicase ERI-6/7 (Fig. 1a). They further demonstrated that a direct repeat flanking *eri-6* facilitated the *trans*-splicing process (Fig. 1a). Remarkably, they also noticed variation of the locus within and between species: a single contiguous gene structure at the *eri-6/7* locus was found in some wild *C. elegans* strains and the *C. briggsae* reference strain AF16. However, the evolutionary history and consequence of the polymorphic variation remained unknown.

Expression quantitative trait loci (eQTL) are genomic loci that explain variation in gene expression across a species^18^. We recently conducted a genome-wide eQTL analysis among 207 wild *C. elegans* strains, using single nucleotide variants (SNVs) as markers^19^ (Extended Data Fig. 1a). Here, we show that the *cis*-acting eQTL of the *eri-6/7* locus is associated with a genomic hotspot enriched for *trans*-acting eQTL of non-conserved genes and pseudogenes, including known ERI-6/7-dependent siRNA targets. We identify structural variation underlying the *eri-6/7* eQTL, including a distinct gene structure and multiple TE remnants. Our results further demonstrate that the insertion of multiple copies of the virus-like DNA transposon, *Polinton*^20,21^, might have caused gene inversion and fission of a single ancestral *eri-6-7* gene. Although some wild strains still possess the single *eri-6-7* gene, other strains such as N2 evolved the *eri-6/7 trans*-splicing mechanism to compensate for the *eri-6* inversion. The direct repeats used for *trans*-splicing originated from the terminal inverted repeats (TIRs) of *Polintons*. The neighboring putative genes *eri-6[e]*, *eri-6[f]*, and *sosi-1* are affected by other *Polinton*-induced structural variants and could have acquired their regulatory functions because of the inversions. Taken together, the *eri-6/7* gene structure polymorphisms and further structural variants at the locus impart sophisticated regulatory effects on the biogenesis of the ERI-6/7 helicase, downstream siRNAs, and the expression of their novel gene targets.

## Results

### Natural variation in *eri-6* underlies differential expression of non-conserved genes and pseudogenes

The genes *eri-6* and *eri-7* are next to each other in an opposite head-to-head orientation at 4.45-4.47 Mb on chromosome I in the N2 reference genome (WS283)^22^ (Fig. 1b). The *eri-6* gene has had a changing transcript annotation in Wormbase^22^ because of a variety of rare splicing events. Presently, it includes six isoforms [*a-f*] that do not all share exons: *eri-6[a-d]* share their first seven exons (hereafter “ERI-6 exons”, which encode the ERI-6 portion of ERI-6/7) and short downstream exons, some of them quite distant; *eri-6[e]* and *eri-6[f]* do not share ERI-6 exons but are transcribed from distinct downstream exons (Fig. 1b). Because the small downstream exons of *eri-6[a-d]* do not contribute many RNA-seq reads, we used the combined expression of *eri-6[a-d]* as a proxy for the total expression of ERI-6 exons (Extended Data Fig. 1b). We investigated the genetic basis of expression variation (eQTL) for ERI-6 exons, *eri-6[e]*, *eri-6[f]*, and other protein-coding genes in *C. elegans* (See Methods and our previous study^19^, Supplementary Tables 1, 2). Here, we focused on eQTL related to the *eri-6/7* locus.

We classified eQTL into local and distant eQTL based on the location of the QTL in the genome relative to its expression targets^19^ (Extended Data Fig. 1a, Supplementary Table 2). At the threshold used (see Methods), we detected local eQTL for expression variation in ERI-6 exons, *eri-6[e]* and *eri-6[f]* (Fig. 1c, Supplementary Table 2). Fine mappings of these local eQTL identified the top candidate variant (I: 4,464,670), a missense mutation (259D>259Y) in the coding region of *eri-6[e]*. Strains with the alternative allele at this site showed significantly lower *eri-6[e]* and *eri-6[f]* expression than strains with the reference allele but higher expression in ERI-6 exons (Fig. 1d). Because *eri-6[e]* was found to repress the expression of ERI-6 exons (Fig. 1a)^17^, it is possible that the alternative non-synonymous allele at the *eri-6[e]* variant could repress the expression of *eri-6[e],* which then would enhance expression of ERI-6 exons.

Expression variation in ERI-6 exons could further affect the production of the ERI-6/7 helicase, the biogenesis of siRNAs in the ERGO-1 pathway, and finally the expression of target genes (Fig. 1a). We found that 13 transcripts of 12 genes across the genome, including four known targets of ERI-6/7-dependent siRNAs^13^, have their distant eQTL (I: 4.3-4.7 Mb) located nearby the *eri-6/7* locus (Fig. 1e, Extended Data Table 1, Supplementary Table 2). Fine mappings of these distant eQTL also identified the I: 4,464,670 *eri-6[e]* variant as the top candidate (Extended Data Table 1). These transcripts showed significantly lower expression in strains with the alternative allele than strains with the reference allele (Extended Data Fig. 1c). Their expression also exhibited negative correlations with ERI-6 exons but positive correlations with *eri-6[e]* expression (Fig. 1f). As mentioned above, pseudogenes and non-conserved genes are among the primary targets of the ERI-6/7-dependent siRNAs^9,13^. Nine of 12 genes are pseudogenes and seven of them lack known orthologs in other species^22^ (Extended Data Table 1). Taken together, all these 12 genes are potential targets of ERI-6/7-dependent siRNAs. Genetic variation at the *eri-6/7* locus functions as a *trans*-acting hotspot to regulate expression of target genes across the genome using the siRNA pathways.

We hypothesized above that the *eri-6[e]* candidate variant could affect the expression of *eri-6[e]*, ERI-6 exons, and potential siRNA targets. However, it was unclear why the variant was also associated with *eri-6[f]* and *sosi-1* expression variation (Fig. 1d). We used CRISPR-Cas9 genome editing to individually introduce the two alleles of the candidate *eri-6[e]* variant into different genetic backgrounds and showed that this variant did not underlie the local eQTL of *eri-6* (Extended Data Fig. 2) nor the distant eQTL of potential targets.

Two of the strains (CB4856 and MY18) in our expression dataset with an alternative allele at the *eri-6[e]* variant were previously found to have *eri-6* and *eri-7* on the same (Crick) strand, similar to the *eri-7* ortholog in the reference genomes of the species *C. briggsae* and *C. brenneri* (Fig. 2)^16,22^. We thus focused on structural variants, which were not included in the eQTL mapping because of the difficulty in characterizing them. We first studied them at the genomic level to uncover the diversity of structural variants, then discovered their transposon origin and finally demonstrated the association of these structural polymorphisms with a diversity of gene expression phenotypes.

**Fig. 2:**
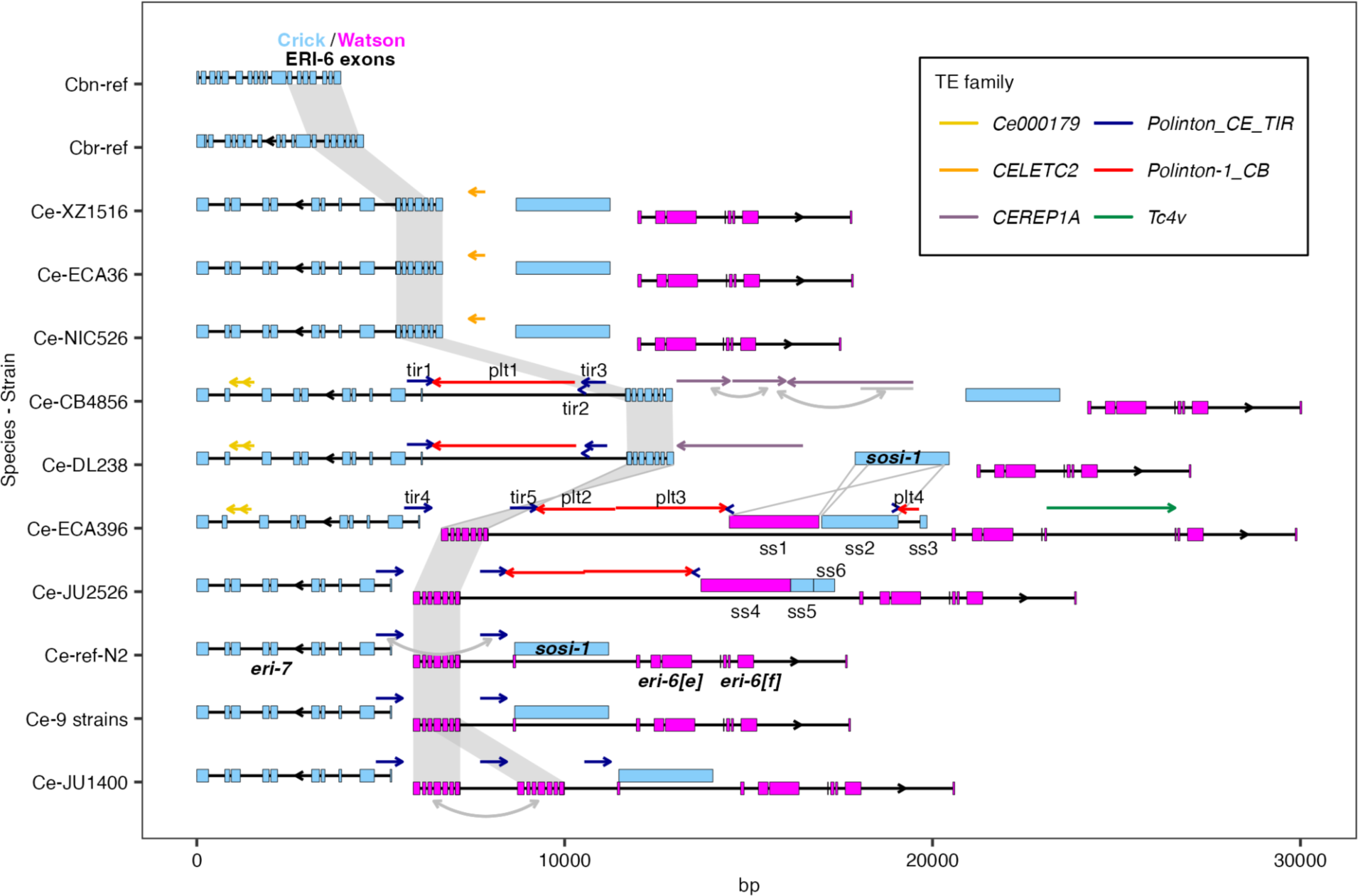
Hyper-variable structural variants and TEs at the *eri-6/7* locus. Graphic illustration of DNA sequence alignment at the *eri-6/7* locus in the 18 *C. elegans* (Ce) strains with genome assemblies. The gene structures of the *C. briggsae* reference (Cbr-ref) *eri-7* and its best match homolog in *C. brenneri* reference (Cbn-ref) are shown on top. The exon structures of the *C. elegans* strains are shown based on the reference N2 genome. Regions with a potential transposon origin are indicated as colored single-headed arrows, with the color indicating the type of transposon and the arrow direction representing their potential coding orientation when inserted. Double-headed arrows indicate duplications. ERI-6 exons are shaded gray. Detailed alignment to the reference of regions with labels “tir1-5” (for terminal inverted repeats), “plt1-4” (for *Polintons*), and “ss1-6” (for *sosi-1*) are shown in Extended Data Fig. 4.

### High diversity of structural variants and TE insertions throughout the *eri-6/7* locus

Long-read genome assemblies of 17 wild *C. elegans* strains are presently available^23–26^, in addition to the reference strain N2. We first performed a multiple pairwise alignment of the *eri-6/7* region among these strains (Fig. 2, Extended Data Fig. 3a)^23–27^. Nine of the 17 strains are approximately identical to the reference strain N2 in this region, with *eri-6* on the Watson strand (pink in figures) and *eri-7* on the Crick strand (blue in figures). Hereafter, the first seven exons of *eri-6* in the N2 reference orientation are called “Watson ERI-6 exons”. The strain JU1400 has a 2.8 kb duplication that includes the Watson ERI-6 exons and one copy of the direct repeats that flank ERI-6 exons (Fig. 2).

The other seven strains harbor a large diversity of deletions, insertions, and inversions compared to the reference genome. The two strains ECA396 and JU2526 have a largely inverted *sosi-1* gene compared to the N2 strain, two different *sosi-1* fragments, and several other insertions (Fig. 2, Extended Data Figs. 3a, 4a). The remaining five strains show inversion of ERI*-* 6 exons compared to the N2 strain (hereafter “Crick ERI*-*6 exons” when in the same orientation as *eri-7*): the strains XZ1516, ECA36, and NIC526 also lack the direct repeats that flank ERI-6 exons and include a ∼1.7 kb insertion between their Crick ERI*-*6 exons and *sosi-1*; the strains CB4856 and DL238 have retained most of the direct repeat sequences and show multiple large insertions with sizes up to ∼8 kb within *eri-7* and surrounding the Crick ERI*-*6 exons (Fig. 2, Extended Data Fig. 3a). The Crick orientation of the ERI-6 exons in these five strains likely represents the ancestral genetic structure at the *eri-6/7* locus, based on the following: 1) *eri-6-7* orthologs in *C. briggsae* and *C. brenneri* show a simple continuous structure on a single strand (Fig. 2); 2) the XZ1516, ECA36, CB4856, and DL238 strains were found to have patterns of ancestral genetic diversity in the *C. elegans* species^28–30^ (Extended Data Fig. 5).

This structural diversity corresponds to an astonishing diversity of polymorphic TEs within the 18 kb locus (Fig. 2, Extended Data Fig. 3a). First, a 435-bp fragment of *CELETC2* (a nonautonomous *Tc2*-related DNA transposon)^22^ resides in the ∼1.7 kb insertion on the right of Crick ERI*-*6 exons in the strains XZ1516, ECA36, and NIC526. Second, two different fragments (354-bp and 299-bp) of the unclassified transposon *Ce000179*^22^ constitute most of the 838-bp insertion within *eri-7* in the strains CB4856, DL238, and ECA396. Third, a full-length *CEREP1A* (a putative nonautonomous 3.4-kb DNA transposon likely using *HAT*-related transposase for propagation)^22^ was found in both the CB4856 and DL238 strains, and the CB4856 strain has two other *CEREP1A* fragments immediately upstream in the opposite orientation. Fourth, the strain ECA396 has a full-length *Tc4v* (a variant class of the DNA transposon *Tc4*)^22,31^ within the first exon of *eri-6[f]*. Fifth, we found multiple TE insertions from a family of autonomous double-stranded DNA transposons derived from viruses, called *Polintons*^20,21^. Four different sizes of *Polinton* remnants were identified at this locus in the strains CB4856, DL238, ECA396, and JU2526 (Fig. 2).

### The direct repeats allowing *eri-6/7 trans*-splicing originate from *Polintons*

*Polintons* (a.k.a. *Mavericks*) were identified across unicellular and multicellular eukaryotes and proposed to transpose through protein-primed self-synthesis^5,20,32^. They code numerous proteins, including two core components, a protein-primed DNA polymerase B (pPolB1) and a retroviral-like integrase (INT), and different capsid proteins^20,21^. The different *Polinton* remnants we found at the *eri-6/7* locus in wild strains are all likely from the pPolB1 end of the *Polinton-1_CB* (named after the *Polintons* in *C. briggsae*, Extended Data Figs. 4b)^22^. In the reference genome of *C. elegans*, three partial copies of *Polinton-1_CB* have been identified at 10.30-10.32 Mb (WBTransposon00000738) and 13.08-13.10 Mb (WBTransposon00000637) on chromosome I and at 17.25-17.27 Mb (WBTransposon00000739) on chromosome X, with lengths ranging from 13.4 to 15.4 kb^22^. We found 744-bp inverted repeats perfectly flanking WBTransposon00000738 (Extended Data Figs. 4b, Supplementary Table 3) and partially flanking the other two *Polintons* in the genome of the reference strain N2. We hypothesized that these inverted repeats were specific terminal inverted repeats (TIRs) of *Polintons* in *C. elegans*. They were previously not regarded as *Polintons* because *C. briggsae Polinton* consensus sequences were used to identify *Polintons* in *C. elegans*. To examine the validity and species specificity of the TIRs, we first identified potential *Polintons* by searching colocalization (within 20 kb) of pPolB1 and INT in the genomes of 18 *C. elegans* and three *C. briggsae* strains (Extended Data Figs. 6). We identified three to nine potential *Polintons* in each *C. elegans* strain and 13 to 15 in each *C. briggsae* strain. Complete or partial sequences of the 744-bp TIRs were flanking 63 of the total 107 *Polintons* in the 18 *C. elegans* strains but none in the three *C. briggsae* strains (Extended Data Figs. 6). We also found colocalization of pPolB1 and the TIR but not INT at 10 loci, including but not limited to the *eri-6/7* locus in *C. elegans* genomes of both N2-like strains and the divergent strains (Extended Data Figs. 6a). Furthermore, all significant NCBI BLAST^33^ results in the query of the TIR sequence are from *C. elegans*. Taken together, the 744-bp TIRs are components of *Polintons* specifically in *C. elegans*, termed *Polinton_CE_TIR*. We distinguish them from the annotated *Caenorhabditis Polinton-1_CB*.

The *Polinton_CE_TIR* sequences are present as direct repeats instead of inverted repeats exclusively at the *eri-6/7* locus in the reference N2, the nine N2-like strains, JU1400, JU2526, and ECA396 (Fig. 2, Extended Data Figs. 6a). In fact, ∼700 bp of the ∼930-bp direct repeats that facilitate *trans*-splicing are exactly *Polinton_CE_TIR* (Extended Data Figs. 3b, Supplementary Table 3). The repeat sequences also include new putative TF binding sites for transcriptional regulation (Extended Data Fig. 3c). Therefore, strains such as the reference N2 use components of *Polintons* to compensate for the disruptive gene inversion that was likely caused by the *Polintons* themselves.

### Multiple *Polinton* copies likely mediated inversions and other structural rearrangements

To evaluate the diversity of this locus using a larger set of strains, we obtained short-read whole-genome sequencing (WGS) data of 550 isotype strains, aligned to the reference N2, representing 1384 wild strains from the *Caenorhabditis* Natural Diversity Resource (CaeNDR, 20220216 release)^34^. We aimed to detect inversions and other structural variants in the species using information of split reads and mapping coverages (See Methods) and relate them to the SNV haplotypes in the region.

We identified diverse structural variants within the *eri-6/7* locus among the 550 wild strains (Extended Data Figs. 3d, 7, Supplementary Table 4): (1) inversions, 93 strains have Crick ERI*-*6 exons and 34 strains have partial inversions of *sosi-1* (INV*sosi-1*) (Extended Data Fig. 7a); (2) *Polinton* insertions, 48 strains likely have partial remnants of the pPolB1 end of the *Polinton-1_CB* (Extended Data Fig. 7a); (3) lack of reference genes (which might result from deletion or maybe an ancestral lack of insertion), 14 strains lack the reference *sosi-1*, *eri-6[e]*, and *eri-6[f]*, whereas two strains only lack *eri-6[e]* and *eri-6[f]* (Extended Data Fig. 7b, d); (4) deletions, 13 strains showed a ∼250-bp deletion mostly spanning the 3’UTR of *eri-6[f]*; (5) duplications, the strain JU1896, might have duplications of *eri-6[e]* and *eri-6[f]*; (6) high heterozygosity in *sosi-1*, 80 strains with the reference *sosi-1* might have a second copy of *sosi-1* beyond the locus, which was also possessed by three of the 14 strains lacking the reference *sosi-1* (Supplementary Table 4).

The short-read data are limited in their ability to detect the full extent of structural variants. However, we observed *Polintons* (*Polinton_CE_TIR* and *Polinton-1_CB*) at multiple sites throughout the *eri-6/7* locus (Extended Data Fig. 7a), especially at flanking regions of ERI*-*6 exons and *sosi-1*. TEs have been associated with chromosomal rearrangements since their first discoveries^1^. Ectopic recombination between TE copies or alternative transposition mechanisms could cause structural variants such as inversions, duplications, or deletions^2^. We reasoned that the inversions of ERI*-*6 exons and *sosi-1* were possibly induced by homologous recombination between the flanking *Polintons* or simply the TIRs.

To understand the evolutionary relationships of the 550 strains at *eri-6/7* and group them, we performed a haplotype network analysis using the 95 SNVs within the locus (Fig. 3). We observed and defined two major groups, “Single *eri-6-7*” and “Reverse-oriented *eri-6,7*”, with 112 and 438 strains, respectively (Fig. 3). As expected, a Crick orientation of ERI-6 exons was detected for all strains in the “Single *eri-6-7*” group, except 17 strains that were clustered with CB4856 and DL238. We hypothesized that all these 17 strains also have the original Crick orientation of ERI*-*6 exons, but with large *Polinton* remnants in between them and *eri-7*: we defined them as “CB4856-like” strains together with the strains DL238 and ECA1186 (Fig. 3). The “Reverse-oriented *eri-6,7*” group of strains includes the reference strain N2 and likely all have Watson ERI*-*6 exons and the direct repeats for *trans*-splicing (Extended Data Figs. 3d, 7b, c, Supplementary Table 4). Most strains in this group are clustered with N2, whereas the strain ECA396 and 19 other strains formed a second cluster based on SNVs and likely all have INV*sosi-1* (Fig. 3). Remnants of *Polinton-1_CB* were found in both groups, but mostly in CB4856-like strains and strains with INV*sosi-1* (Fig. 3). Strains with deletion polymorphisms in *eri-6[e]*, *eri-6[f]*, and *sosi-1* formed two clusters exclusively in the “Single *eri-6-7*” group (Fig. 3). It is challenging to associate these structural variants with *Polintons* or other TEs. Nevertheless, these deletions and duplications might also affect expression of *eri-6/7* and siRNA pathways.

**Fig. 3:**
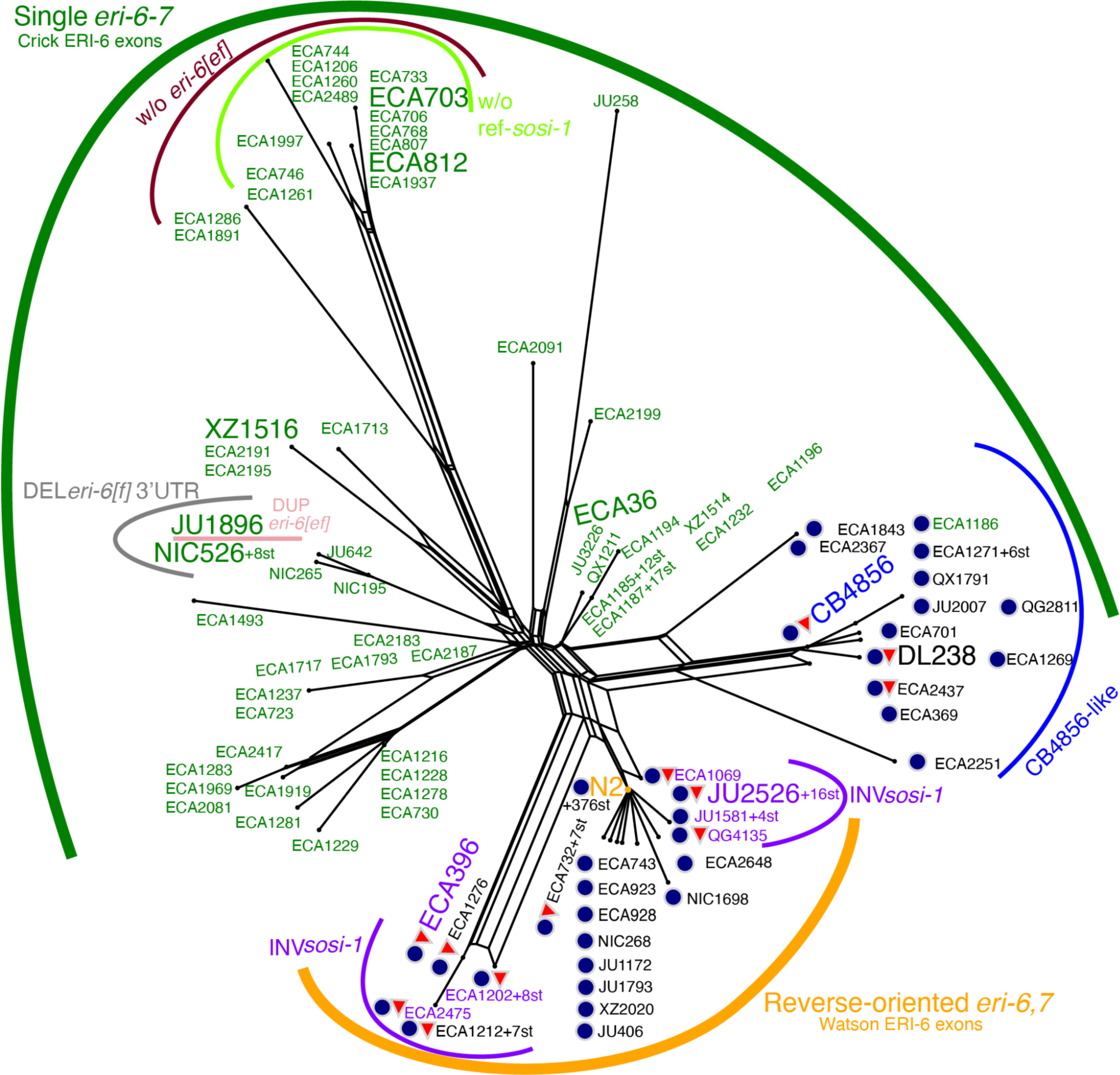
Haplotype network with clustered strains sharing structural variation. Neighbor-joining net depicting 550 strains based on 95 SNVs within the *eri-6/7* locus. Two major groups, “Single *eri-6-7*” and “Reverse-oriented *eri-6,7*”, were defined based on orientation of ERI-6 exons and denoted with dark green and orange curves. Subgroups with other structural variations were indicated using thin curves and labels (“w/o” for deletions or no insertions, “DEL” for deletions, and “DUP” for duplications). Strain names are colored in green and purple for detection of Crick ERI-6 exons and inversion of *sosi-1* (INV*sosi-1*), respectively, using short-read WGS data in Extended Data Fig. 7a. Dark blue circles and red triangles next to strain names represent strains with *Polinton_CE_TIR* (TIRs only) and *Polinton-1_CB* (TIRs excluded) insertions, respectively, based on Extended Data Figs. 3d, 7 and manual inspection of genome alignments. Some strains (st) share all alleles of the 95 SNVs and all detected structural variations are collapsed to only show a representative strain followed by the number of strains with this *eri-6/7* haplotype (e.g. “N2 +376st”). Trapezoidal junctions indicate that some recombination occurred within the locus.

### *Cis*-and *trans*-effects of *Polinton*-induced structural variants on gene expression

Among the 550 wild *C. elegans* strains, ∼20% likely have a single “classical” *eri-6-7* gene to encode the ERI-6/7 protein, as in *C. briggsae* and *C. brenneri*. The remaining ∼80% strains make a fused *eri-6/7* mRNA by some amount of *trans*-splicing between the pre-mRNAs of the Watson ERI-6 exons and *eri-7* as in the reference strain N2. Though *trans*-splicing compensates inversion of ERI-6 exons to continue ERI-6/7 production, Fischer and Ruvkun could not consider whether the reverse-oriented *eri-6/7* gene structure might represent a hypomorphic form of the locus compared to the ancestral, compact gene. We thus turned our focus back to gene expression consequences of structural variants, which could affect expression at two levels: the expression abundances of different exons and their splicing.

We first examined local regulatory effects at the *eri-6/7* and *sosi-1* locus, starting with diversity among strains having the Crick ERI-6 exons and *eri-7*. The strains with a potential compact *eri-6-7* gene (green box color in Fig. 4a) expressed both parts of the gene at similar levels, as expected, and expressed low levels of *eri-6[e]*, *eri-6[f]*, and *sosi-1*. The exception in this group is the two strains ECA703 and ECA812, which do not have *eri-6[e]*, *eri-6[f]*, and *sosi-1* and showed low expression in ERI-6 exons and ref-*eri-7* (mRNA sequences of *eri-6[a-d]* and *eri-7* in the N2 reference, respectively) (Figs. 3, 4a, Supplementary Table 4). Because *eri-6[e]*, *eri-6[f]*, and *sosi-1* were found to repress *eri-6/7* expression in the reference strain N2^17^, their putative deletions could cause elevated expression of ERI-6 exons and *eri-7*. These observations suggest that other linked genetic variation at the locus reduces expression of ERI-6 exons and ref-*eri-7* or that *eri-6[e]*, *eri-6[f]*, and *sosi-1* function differently in strains of the “Single *eri-6-7*” group. The strain JU1896, which likely has a duplication in *eri-6[e]* and *eri-6[f]* showed higher expression in both (Figs. 3, 4a, Extended Data Fig. 7d). The subgroup of CB4856-like strains (blue color), with large *Polinton* remnants between ERI-6 exons and the downstream ERI-7 exons (Fig. 2), exhibited significantly elevated expression in ERI-6 exons and significantly decreased expression in ref-*eri-7*: the large intronic insertion likely affects transcription of the downstream exons, *i.e.*, *eri-7*.

**Fig. 4:**
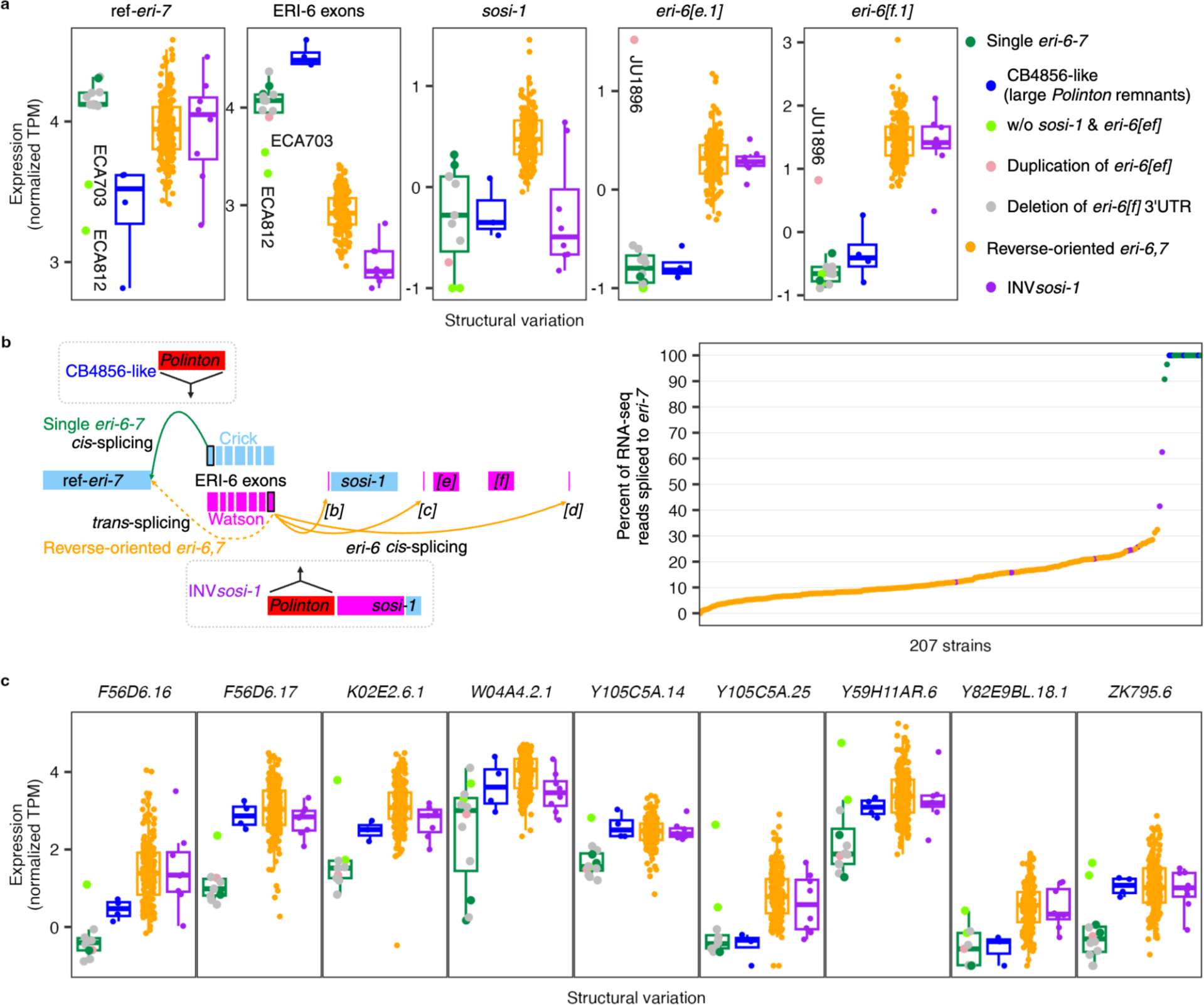
Structural variations at the *eri-6/7* locus regulate genes in *cis* and *trans*. **a**,**c**, Tukey box plots showing expression (-*log*_2_(normalized TPM+0.5)) variation of five transcripts at the *eri-6/7* locus (**a**) and nine transcripts (**c**) across the genome that include known targets of siRNAs requiring the ERI-6/7 helicase, among strains with major and minor structural variations (SVs) within the locus. Each data point represents a strain, color-coded by SVs. Each box is colored by major SVs. Box edges denote the 25^th^ and 75^th^ quantiles of the data; and whiskers represent 1.5× the interquartile range. Statistical pairwise comparison results using two-sided Wilcoxon tests and Bonferroni corrections were presented in Supplementary Table 5. **b**, Percent of spanning RNA-seq reads at the end of the last (seventh) ERI-6 exon that were spliced to *eri-7* when mapped to the reference genome, for 207 strains. Each point represents one strain and is colored by SVs. Graphic illustration of structural variation within the *eri-6/7* locus was created using BioRender.

The second large group of strains, those in the “Reverse-oriented *eri-6,7*” group (orange and purple colors), showed significantly lower expression in ERI-6 exons and significantly higher expression in *eri-6[e]*, *eri-6[f]*, and *sosi-1* than strains in the “Single *eri-6-7*” group (Fig. 4a, Supplementary Table 5). The lower expression ERI-6 exons might be the result of either enhancer/promoter rearrangement or deficiencies in splicing or poly-A tail formation making the mRNA less stable. By contrast, these strains exhibited a similar level of expression of the ref-*eri-7* to the “Single *eri-6-7*” group. The subgroup of strains with INV*sosi-1* (purple color) showed significantly lower expression in both *sosi-1* and ERI-6 exons than other strains in the “Reverse-oriented *eri-6,7*” group. Those strains with genome assemblies show large *Polinton* remnants upstream of *sosi-1* (Fig. 2), which could explain the lower expression of *sosi-1* and perhaps render mRNAs of ERI-6 exons unstable. In summary, the diverse structural variations correlate with their expected effect on the *eri-6/7* locus between and within the two large structural variant groups.

Different splicing mechanisms between the two groups further alter the efficiency of the ERI-6/7-dependent siRNA pathways. In strains with a single *eri-6-7* gene, the ERI-6/7 protein is produced through standard transcription and translation. In contrast, strains with reverse-oriented *eri-6,7* perform separate transcription of pre-mRNAs in opposite orientation and *trans*-splicing^16^, which could reduce the efficiency of ERI-6/7 production. We analyzed spanning reads in the RNA-seq data of 207 strains to compare their splicing efficiency between the seventh exon of *eri-6* and the first of *eri-7* (Fig. 4b). In strains with a single *eri-6-7* gene, most split RNA-seq reads at the end of the Crick ERI-6 exons should have their chimeric alignment to ERI-7 exons through *cis*-splicing. In strains with the Watson ERI-6 exons, split RNA-seq reads at the end of ERI-6 exons could splice to downstream exons for *eri-6[b-d]* or partially map to ERI-7 exons because of *trans*-spliced *eri-6/7* mRNAs^16^ (Fig. 4b). Indeed, among the 207 strains in our RNA-seq dataset, all 16 strains with a single *eri-6-7* gene showed higher than 90% and mostly 100% splicing between ERI-6 and ERI-7 exons. Instead, the 183 strains with reverse-oriented *eri-6,7* but not INV*sosi-1* showed a median of 10% and a maximum of 32% *trans*-splicing (Fig. 4b). In conclusion, the evolutionary inversion of *eri-6* does affect the synthesis of full-length *eri-6/7* mRNA.

Together, the expression level of ERI-6 and ERI-7 exons and their splicing rate alter the biogenesis of the helicase ERI-6/7. Strains with a single *eri-6-7* gene but no extra insertions or deletions might generate the most abundant ERI-6/7 because of their high expression in ERI-6/7 exons and mostly 100% *cis*-splicing (Fig. 4a, b). The reverse-oriented *eri-6/7* gene structure represents a hypomorphic form of the locus, because strains in this group showed decreased expression of ERI-6 exons and low splicing rate between ERI-6/7 exons (Fig. 4a, b), which likely causes reduced ERI-6/7 protein.

The structural variants showed various local effects on gene expression but their influences likely extend beyond the locus because of the pivotal role of ERI-6/7 in *C. elegans* endogenous siRNA pathways (Fig. 1a). Differences in ERI-6/7 abundances will affect the generation of ERGO-1 dependent siRNAs and repression of their target genes. Among the putative targets of ERI-6/7-dependent siRNAs from our eQTL analysis, we observed significantly lower expression in strains in the “Single *eri-6-7*” group than strains in the “Reverse-oriented *eri-6,7*” group (Fig. 4c, Supplementary Table 5). We also found potential effects of structural variants in the CB4856-like strains on target expression variation within the “Single *eri-6-7*” group. Altogether, these results demonstrate that diverse structural variants at the *eri-6/7* locus altered *C. elegans* endogenous siRNA pathways from the production of the ERI-6/7 helicase to the expression of target genes.

## Discussion

### Evolutionary genomic history of the *eri-6/7* locus driven by *Polintons*

Most strains with a single *eri-6-7* gene were isolated from the Hawaiian Islands or the Pacific region, where the highest known genetic diversity in the *C. elegans* species is found, (Fig. 3, Extended Data Figs. 5, 8), which likely reflects the retention of ancestral diversity^28–30^. Strains with an inversion of ERI-6 exons, however, are more widely distributed over the world and predominant in Europe. This set of strains show reduced genetic diversity at the locus, in agreement with an evolutionary-derived inversion of ERI-6 exons from the Crick to the Watson strand within the species (Extended Data Figs. 5, 8)^30^.

We thus favor the following scenario of evolution at the *eri-6/7* locus (Fig. 5). The *eri-6/7* gene was ancestrally coded as a single gene as in *C. briggsae* and *C. brenneri*, without *Polinton* insertions. The lack of *eri-6/7* homolog in *C. inopinata*^14^ prevents us from using it as a closer outgroup. The ancestor of all *C. elegans* strains likely conserved the compact single *eri-6-7* gene structure as in *C. briggsae* and *C. brenneri*. Some strains, such as XZ1516, likely kept this ancestral single *eri-6-7* gene with no trace of *Polintons* (Figs. 2, 5). Alternatively, in these strains, the *Polintons* were fully eliminated from the *eri-6/7* locus, yet the parsimonious explanation is that *Polintons* invaded the locus after the speciation of *C. elegans*.

**Fig. 5:**
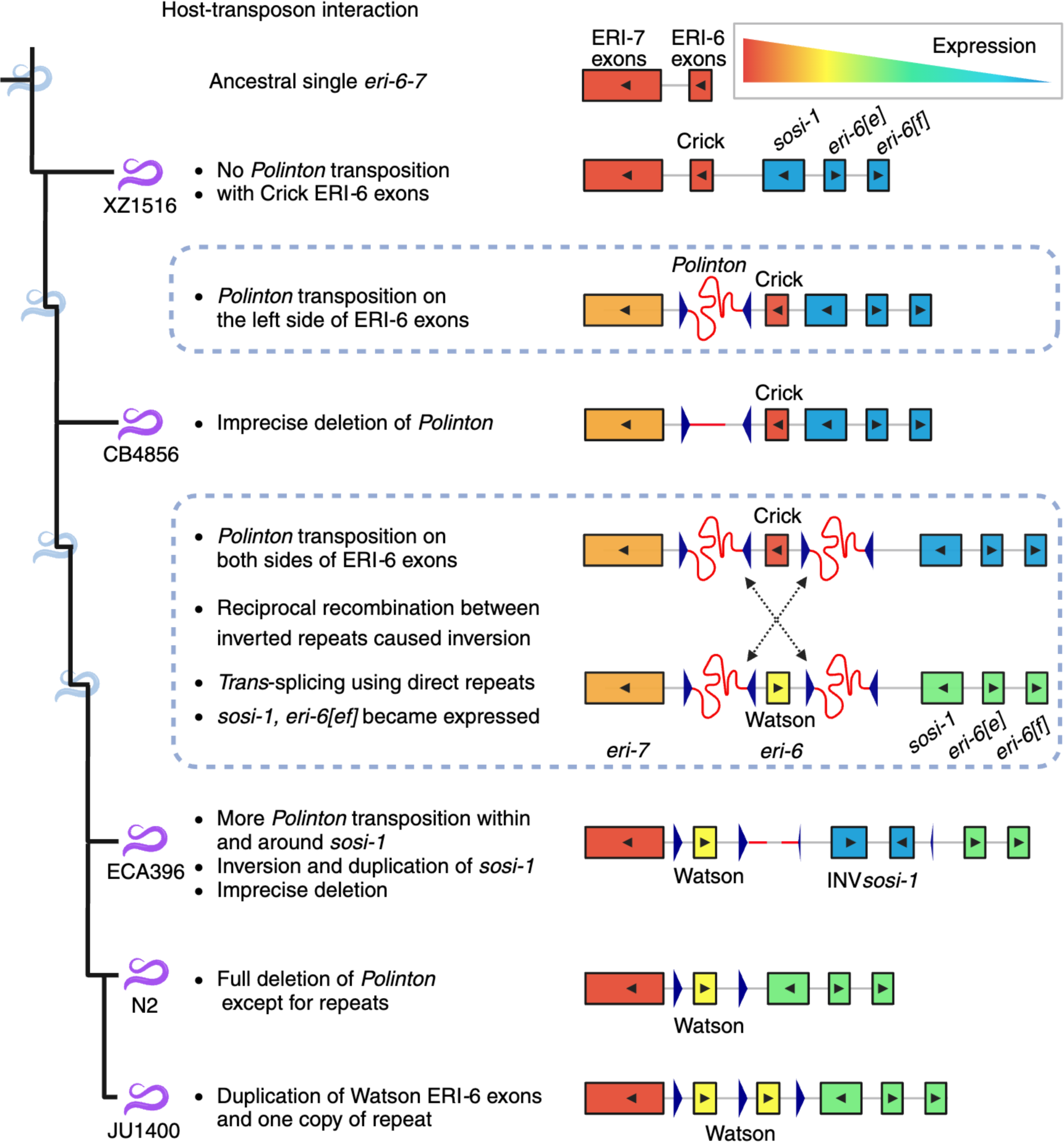
Possible scenario for evolution at the *eri-6/7* locus with *Polintons*. Purple and light blue worms on the tree represent nodes with or without actual strains, respectively, to the best of our knowledge. Rectangles for different segments of *eri-6/7* were filled with gradient colors to indicate expression level across segments and branches on the tree. Black triangles inside rectangles represent orientation of gene segments. Dark blue triangles represent repeats. Red curved lines indicate *Polintons* other than the repeats. Created using BioRender.

We found *Polinton* remnants in the genome of every *C. elegans* strain with available WGS data at CaeNDR (Extended Data Fig. 6). At some time during the evolutionary history of *C. elegans*, a *Polinton* copy transposed, likely from another location in the genome or through horizontal transfer, and interrupted the *eri-6/7* gene with a large insertion on the left side of ERI-6 exons. No strain in our dataset retains a full *Polinton* at the locus, thus this *Polinton* was either a partial copy when it transposed or subsequently became largely deleted. In strains such as CB4856, the still large *Polinton* remnants (∼5 kb in CB4856) appear to impair *eri-7* transcription (Figs. 4a, 5).

Further *Polintons* insertions occurred in the vicinity, including perhaps to the right side of ERI-6 exons (Figs. 2, 5). The occurrence of several *Polinton* copies at the same locus may have favored ectopic recombination between inverted sequences and the ERI-6 exon inversion (Fig. 5). Surviving descendants of this inversion, such as the ECA396 and N2 strains, use repeats from *Polintons* for *trans*-splicing and thus maintain a hypomorphic *eri-6/7* function (Figs. 4c, 5). Meanwhile, the inversion activated *eri-6[e]* and *eri-6[f]*, which were barely expressed in most strains with a single *eri-6-7* gene, at least in the tested conditions (Figs. 4a, 5). Ancestors of the reference strain N2 eliminated other *Polinton* fragments from the locus, except for the direct repeats that are necessary for *trans*-splicing. Strains such as JU1400 evolved a duplication of the Watson ERI-6 exons and one copy of the direct repeat, which may increase the number of correctly spliced *eri-6/7* transcripts (Figs. 2, 5). *Polintons* might have caused more structural variations such as INV*sosi-1* (Figs. 2, 5, Extended Data Fig. 7).

The actual evolutionary process within this locus must be more complex than the model proposed above. The *Polinton* insertions could have occurred through sudden bursts of transposition instead of gradually. Sudden environmental stress might have caused the high transposition rate of *Polintons* and the other four TEs (Fig. 2). Overall, the large number of transposon insertions at this locus regulating small RNA pools and thereby transposons support the hypothesis of a presumed battle between TE insertions and genomic rearrangement to preserve ERI-6/7 function to combat further TE activity. Only through further investigations of gene expression and TE positions in *de novo* assemblies will we learn more about the broad evolutionary significance of this type of battle.

### Phenotypic effect of the structural variation at *eri-6/7* on siRNA pathways and their targets

With the ERGO-1 Argonaute, the ERI-6/7 helicase is required for production of endogenous primary 26G siRNAs by non-canonical Dicer processing of target mRNAs^13^. Secondary siRNAs are produced by an amplification machinery, for which different pools of primary siRNAs compete^15,35^, including endo-siRNAs dependent on Argonautes ERGO-1 and ALG-3/4, the genomically encoded piRNAs, and the siRNAs derived from exogenous double-stranded RNAs^13,16,36–38^. Depending on the genomic and environmental contexts, genetic variation favoring one or the other primary siRNA pathway could have been selected^39–42^. Research in mammals has shown the importance of dosage of the orthologous MOV10 helicase on retrovirus silencing^43^. We showed here that natural structural variants at the *eri-6/7* locus were a major driver of variation in ERGO-1 pathway activity and mRNA levels of its downstream regulated targets. Two events, likely driven by *Polintons*, lowered ERI-6/7 pathway activity, and increased piRNA-dependent and exogenous RNAi pathways: (1) the initial insertion of a *Polinton* within the *eri-6/7* gene and (2) the inversion of ERI-6 exons. Other events might have acted in the reverse direction: the deletion of most of the intervening *Polintons*, the retention of direct repeats used in *trans*-splicing and, in the strain JU1400, the duplication of the inverted ERI-6 exons. Because ERI-6/7-dependent siRNAs primarily target retrotransposons and unconserved, duplicated genes, with few introns, potentially of viral origins^9,13^, the insertion of the *Polintons* and the resulting inversion could have at least transiently increased expression of novel genes and retrotransposons, while repressing exogenous dsRNAs.

However, it is unclear what the effect might have been on *Polintons* themselves. Since their recent discovery in *C. elegans,* their possible regulation by small RNAs remains to be studied. The DNA polymerase of *Polintons* might be an ancient target of ERI-6/7-dependent siRNAs, because the gene *E01G4.5*, a known target of ERI-6/7-dependent siRNAs in *C. elegans*, encodes a protein that has homology to viral DNA polymerases^9,13^. *Polintons* might also bring novel genes within them^5^, which are potential targets of the ERGO-1 or piRNA pathways. The genes *sosi-1*, *eri-6[e]*, and *eri-6[f]* are absent at the *eri-6/7* locus in a subset of Hawaiian strains showing the most divergent *eri-6/7* region based on SNVs (Fig. 3). It is tempting to suggest that they appeared at this locus during the evolution of the species. The *eri-6[f]* exons are highly similar to another locus in the genome^17^. The gene *sosi-1* keeps additional copies in some wild strains and is a distant paralog of *eri-7* and other helicases in its C-terminal part. Further research can test whether *sosi-1*, *eri-6[e]*, and *eri-6[f]* have been carried by a *Polinton* transposon. Similarly, the mode of duplication of the ERI-6/7 targets remains to be investigated.

Detailed genetic studies in the N2 reference strain have uncovered intricate regulatory interactions at the *eri-6/7/sosi-1* locus and between this locus and the splicing machinery. First, in the N2 strain, in part through matching piRNAs, *eri-6[e]*, *eri-6[f]*, and *sosi-1* are strong ERI-6/7-independent siRNA targets^17^. Their downregulation by MUT-16-dependent siRNAs enables *eri-6/7* expression, perhaps by spreading chromatin marks^17^. This regulation has been proposed to act as a negative feedback loop balancing ERGO-1 dependent secondary siRNAs and other secondary siRNA classes. Second, the use of the *Polinton* repeats as *trans*-splicing signal partially rescues the production of ERI-6/7. This peculiar mechanism of *eri-6/7 trans*-splicing was proposed to act as a compensatory sensor of the splicing machinery, enabling more exogenous RNAi when an overwhelmed splicing machinery increases endo-siRNA production on poorly spliced genes^44^. It remains unclear whether these seemingly intricate effects on siRNA pools in the N2 reference strain are an evolutionary leftover of transposon-driven structural variation at the locus. We hypothesize that across the evolutionary history of *C. elegans*, different siRNA pools may have been successively favored by natural selection. Alternatively, successive structural variants could have endowed the *eri-6/7* locus with physiological regulatory loops used in balancing the different siRNA classes downstream of environmental and organismal inputs.

To conclude, our work dissected a distant eQTL hotspot and identified diverse TEs and structural variations within the *eri-6/7* locus underlying variation in *C. elegans* endogenous siRNA pathways. This locus appears to have been the target of a large number of TE insertions including multiple copies of the otherwise rare *Polinton* transposon, which may have caused high genetic diversity at the locus through genome rearrangements. Some *C. elegans* strains evolved an odd *trans*-splicing mechanism to maintain hypomorphic function of the locus, using *Polinton* TIRs that came to form direct repeats. The remarkable interactions between hosts and TEs play a major role in genome rearrangements and the regulation of gene expression.

## Methods

### Genomic and transcriptomic data

We obtained the reference genomes of *C. elegans* (N2) and *C. briggsae* (AF16), the GTF files of *C. elegans*, *C. briggsae*, and *C. brenneri*, from WormBase (WS283)^22^; the *de novo* assemblies of 17 wild *C. elegans* strains (CB4856, DL226, DL238, ECA36, ECA396, EG4725, JU310, JU1395, JU1400, JU2526, JU2600, MY2147, MY2693, NIC2, NIC526, QX1794, XZ1516) and two wild *C. briggsae* strains (QX1410, VX34) from the NCBI Sequence Read Archive (SRA projects PRJNA523481, PRJNA622250, PRJNA692613, PRJNA784955, and PRJNA819174)^23–27^, the alignment of whole-genome sequence data in the BAM format of 550 wild *C. elegans* strains, the soft- and hard-filtered isotype VCF from the *Caenorhabditis* Natural Diversity Resource (CaeNDR, 20220216 release)^34^; the Illumina RNA-seq FASTQ files of 608 samples of 207 wild *C. elegans* strains from the NCBI SRA (projects PRJNA669810)^19^.

### RNA-seq mapping and eQTL analysis

To put transcriptomic data on the same page with the genomic data, we re-mapped RNA-seq reads using the *C. elegans* reference genome (WS283), the GTF file (WS283), and the pipeline *PEmRNA-seq-nf* (v1.0) (https://github.com/AndersenLab/PEmRNA-seq-nf). Then, we selected reliably expressed transcripts, filtered outlier samples, and normalized expression abundance across samples using the R scripts *counts5strains10.R*, *nonDivergent_clustered.R*, and *norm_transcript_gwas.R* (https://github.com/AndersenLab/WI-Ce-eQTL/tree/main/scripts), respectively, as previously described^19^. In summary, we collected reliable expression abundance for 23,349 transcripts of 16,172 genes (15,449 protein-coding genes and 723 pseudogenes) from 560 samples of 207 strains. We also used *STAR* (v2.7.5) ^45^ to identify chimeric RNA-seq reads in the 560 samples.

We further used our recently developed GWAS mapping pipeline, *Nemascan*^46^, to identify eQTL for the 23,349 transcript expression traits (Supplementary Table 1), following the steps outlined previously^19^. Briefly, we randomly selected 200 traits and permuted each of them 200 times. For each of the 40,000 permuted traits, we used the leave-one-chromosome-out (LOCO) approach and the INBRED approach in the *GCTA* software (v1.93.2)^47,48^, and calculated the eigen-decomposition significance (EIGEN) threshold as *-log_10_(0.05/N_test_)* to identify QTL.

We determined the 5% false discovery rate (FDR) significance threshold for LOCO and INBRED, respectively, by calculating the 95^th^ percentile of the significance of all detected QTL above using each approach. We then performed GWAS mapping on all 23,349 traits using LOCO and INBRED approaches and identified eQTL that passed their respective 5% FDR thresholds. Overall, we detected 10,291 eQTL for 5668 transcript expression traits, with 4899 eQTL for 4254 traits in LOCO, 5392 eQTL for 4700 traits in INBRED (Supplementary Table 2). Fine-mappings were further performed on each eQTL using *Nemascan*.

We classified eQTL as local (within 2 Mb surrounding the transcript) or distant (non-local). For distant eQTL located outside of the common hyper-divergent regions among the 207 strains^19,25^, we identified hotspot regions enriched with distant eQTL for LOCO and INBRED results, respectively^19^.

The genomic region harboring the *eri-6/7* locus at 21 cM on chromosome I was identified as a distant eQTL hotspot in both LOCO and INBRED in this study and in our previous study^19^.

### DNA alignment

We aligned each of the 17 *de novo* PacBio assemblies of wild *C. elegans* strains to the N2 reference genome using *MUMmer* (v3.1)^49^ and extracted sequences that were aligned to the N2 *eri-6/7* locus using *BEDTools* (v2.29.2)^50^. Then, we performed pairwise alignments among these sequences and to the *eri-6/7* N2 reference sequence using *Unipro UGENE* (v.47.0)^51^. Large insertions (>50 bp) in the wild strains to the reference were blasted in WormBase^22^ to identify potential transposon origins.

### Scan for *Polinton* and TIRs in genome assemblies

We obtained the amino acid sequences of pPolB1 and INT in *C. briggsae Polinton-1* (WBTransposon00000832)^22^ using *ORFfinder* (https://www.ncbi.nlm.nih.gov/orffinder/) and the 744 bp DNA sequence for the TIRs from 10,302,516 to 10,303,259 bp on chromosome I in the *C. elegans* (N2) reference genome. We searched for the *Polinton* and TIRs sequences in the 21 genome assemblies using *tblastn* and *blastn* in BLAST (v2.14.0)^52^, respectively. We filtered the results by a maximum *e*-value of 0.001 and a minimum bitscore of 50^32^. We merged pPolB1, INT, and TIR hits within 4 kb, 2 kb, and 2 kb, respectively, with consideration of strandedness. *Polinton* insertions were identified by the presence of both pPolB1 and INT within 20 kb.

We also searched for *sosi-1* outside of the *eri-6/7* locus in the genome assemblies using DNA sequence of *sosi-1* in the reference and found an additional copy in the strains JU2526, ECA396, XZ1516, and JU1400, and two additional copies in the strains ECA36 and QX1794 in their PacBio genome assemblies. Genomic locations surrounding these additional copies in the six strains correspond to ∼0.31 Mb on the chromosome III in the reference N2 genome. The additional copies of *sosi-1* outside the *eri-6/7* locus in the six strains share most alleles compared to the *sosi-1* within the *eri-6/7* locus.

### Identification of SVs using short-read WGS data

We extracted information of split reads mapped to the reference *eri-6/7* locus (I: 4,451,194 - 4,469,460 bp) and with a minimum quality score equal of 20 from the BAM files of the 550 wild *C. elegans* strains. 1): To identify potential inversions in the *eri-6/7* locus, we first selected split reads with both the primary and chimeric alignments mapped to this region but to different strands. We assigned the primary and chimeric alignment positions of each split read into 200-bp bins and required at least four reads that had the primary and chimeric alignments in the same pair of bins for a relatively reliable inversion event in each strain. We focused on inversions spanning at least three bins and found in more than 10 strains. 2): To identify potential sites of *Polinton* remnants, we selected the split reads outside of the direct repeats at the *eri-6/7* locus and with the chimeric alignment mapped to *Polinton (Polinton-1_CB*, WBTransposon00000738) and its surrounding *Polinton_CE_TIR* on chromosome I from 10,302,516 to 10,319,657 bp. At least two reads were required. The primary alignment of these reads indicated the potential sites of *Polinton* remnants in the *eri-6/7* locus in wild strains.

Furthermore, we counted the coverage per bp in the *eri-6/7* locus for each short-read WGS BAM file using *BEDTools* (v2.29.2)^50^. We calculated the percentage of the coverage at each bp to the mean coverage within the *eri-6/7* locus in each strain. Then, we performed a sliding window analysis with a 200-bp window size and a 100-bp step size for each strain. A 173-bp tandem repeat region from 4,465,414 to 4,465,586 bp on chromosome I was masked in the results.

To identify additional copies and haplotypes of *sosi-1* among the 550 wild strains, we focused on 93 variants of the 101 SNVs tagged “high heterozygosity” within the *sosi-1* region in the soft-filtered isotype VCF (CaeNDR, 20220216 release)^34^. We used the following threshold to define *sosi-1* haplotype and copy numbers among the 550 strains: 449 strains show homozygous reference alleles at all 93 SNVs (except one strain at 92 SNVs), indicating they only have the reference haplotype *sosi-1*; 80 strains show heterozygous alleles at more than 60 SNVs, indicating two copies of *sosi-1* with divergent haplotypes; three strains have homozygous alternative alleles at more than 90 SNVs, indicating missing of the reference *sosi-1* in the *eri-6/7* locus and the existence of the alternative *sosi-1* copy; 11 strains show undetected genotype at 60 to 93 SNVs and extreme low coverages in *sosi-1* (Extended Data Fig. 7d), indicating they may lack *sosi-1* in the genomes; the *sosi-1* haplotype and copy number of the remaining seven strains are unclear as they have numbers of homozygous and homozygous alleles in between the above threshold (Supplementary Table 4).

### Genetic relatedness

Genetic variation data across the genome among the 550 *C. elegans* strains were extracted from the hard-filtered VCF above using *BCFtools* (v.1.9)^53^. These variants were pruned to the 1,199,944 biallelic SNVs without missing genotypes. We converted this pruned VCF file to a PHYLIP file using the *vcf2phylip.py* script^54^. The unrooted neighbor-joining tree was made using the R packages *phangorn* (v2.5.5)^55^ and *ggtree* (v1.14.6)^56^.

A second PHYLIP file was built by the same method above but only with 95 SNVs within the *eri-6/7* locus. A haplotype network was generated using this PHYLIP file and *SplitsTree CE* (v6.1.16)^57^.

## Acknowledgements

This research was supported by the National Science Foundation (1764421) and Simons Foundation (597491) Research Center for Mathematics of Complex Biological Systems and a Human Frontiers Research Program grant (RGP0001/2019) to E.C.A. G.Z. is supported by a grant from the Fondation pour la Recherche Médicale (ARF202209015859). M.-A.F. is supported by the Centre National de la Recherche Scientifique and a grant from the Agence Nationale de la Recherche (ANR-19-CE12-0025). We like to thank Emily Koury for the efforts in CRISPR-Cas9 genome editing. We like to thank Nicolas D. Moya for suggestions on useful bioinformatic tools. We would like to thank members of the Andersen Lab for helpful comments on the manuscript. We also like to thank CaeNDR (supported by NSF Capacity Grant 2224885 to E.C.A.) and WormBase for which these analyses would not have been possible.

## Author contributions

G.Z., M.-A.F., and E.C.A. conceived of the study. G.Z. analyzed the data. G.Z., M.-A.F., and E.C.A. wrote the manuscript.

## Data and code availability

The datasets and code for generating all figures can be found at https://github.com/AndersenLab/Ce-eri-67

## Supplementary information

Description of Additional Supplementary Files

Supplementary Table 1

23,349 GWAS gene expression traits

Supplementary Table 2

eQTL summary

Supplementary Table 3

Sequences alignment of *Polinton_CE_TIR* and the direct repeat

Supplementary Table 4

Structural variants at the *eri-6/7* locus of 550 strains

Supplementary Table 5

Statistical pairwise comparison results in Fig. 4a, c

## Extended data figures and tables

**Extended Data Table 1.**
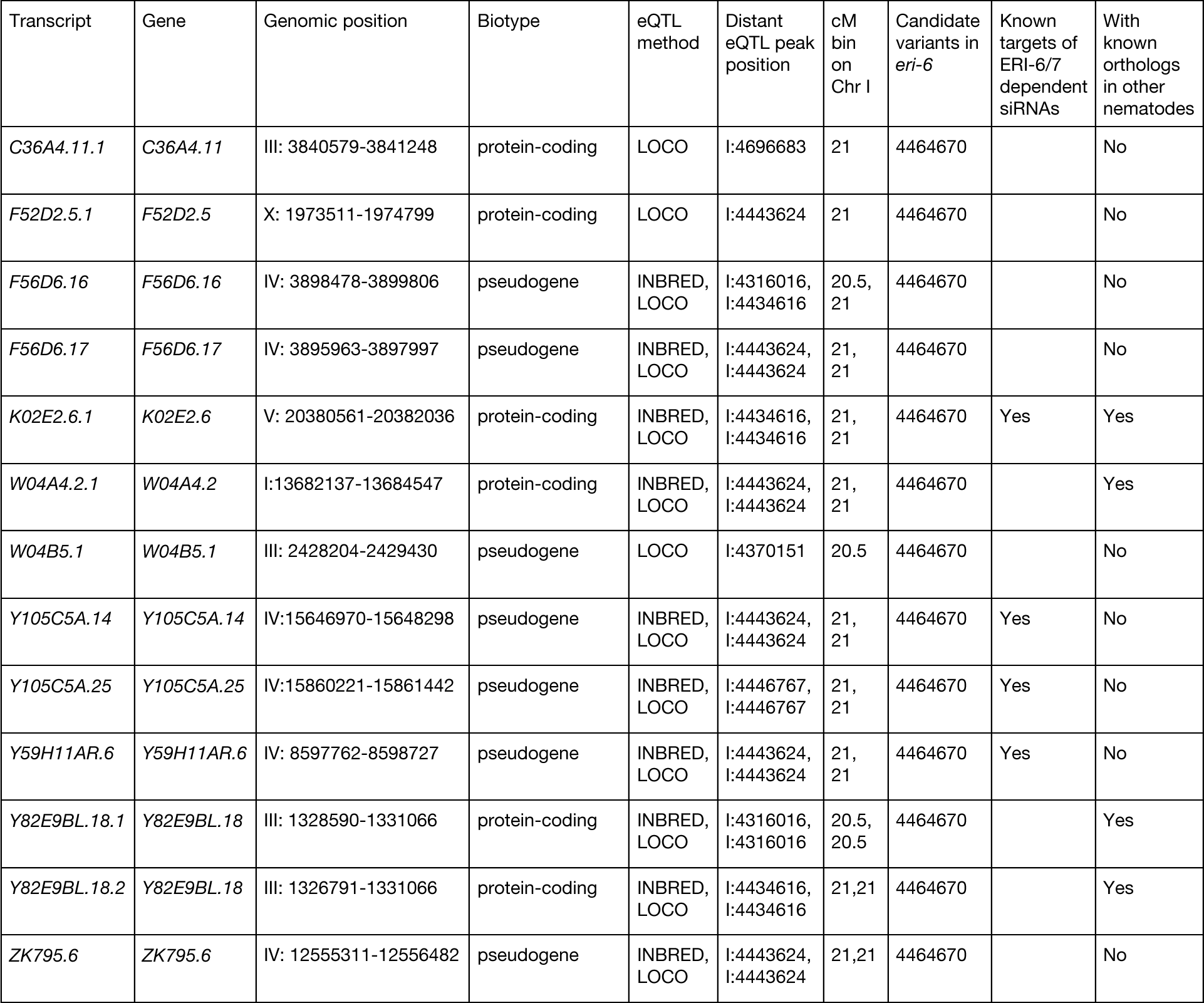
Transcript expression traits associated with *eri-6*. In Fig. 1e, we showed Manhattan plots for the ten traits identified with distant eQTL using both INBRED and LOCO methods. In Extended Data Fig. 1c, we showed the phenotype by genotype plots for all of ten gene expression traits at the top candidate variant. Because of the negative correlation in expression between the two transcripts *Y82E9BL.18.1* and *Y82E9BL.18.2* of the gene *Y82E9BL.18*, only one of these transcripts was depicted in Fig. 4c for clarity purposes.

| Transcript | Gene | Genomic position | Biotype | eQTL method | Distant eQTL peak position | cM bin on Chr I | Candidate variants in *eri-6* | Known targets of ERI-6/7 dependent siRNAs | With known orthologs in other nematodes |
| --- | --- | --- | --- | --- | --- | --- | --- | --- | --- |
| *C36A4.11.1* | *C36A4.11* | III: 3840579-3841248 | protein-coding | LOCO | I:4696683 | 21 | 4464670 |  | No |
| *F52D2.5.1* | *F52D2.5* | X: 1973511-1974799 | protein-coding | LOCO | I:4443624 | 21 | 4464670 |  | No |
| *F56D6.16* | *F56D6.16* | IV: 3898478-3899806 | pseudogene | INBRED, LOCO | I:4316016, I:4434616 | 20.5, 21 | 4464670 |  | No |
| *F56D6.17* | *F56D6.17* | IV: 3895963-3897997 | pseudogene | INBRED, LOCO | I:4443624, I:4443624 | 21, 21 | 4464670 |  | No |
| *K02E2.6.1* | *K02E2.6* | V: 20380561-20382036 | protein-coding | INBRED, LOCO | I:4434616, I:4434616 | 21, 21 | 4464670 | Yes | Yes |
| *W04A4.2.1* | *W04A4.2* | I:13682137-13684547 | protein-coding | INBRED, LOCO | I:4443624, I:4443624 | 21, 21 | 4464670 |  | Yes |
| *W04B5.1* | *W04B5.1* | III: 2428204-2429430 | pseudogene | LOCO | I:4370151 | 20.5 | 4464670 |  | No |
| *Y105C5A.14* | *Y105C5A.14* | IV:15646970-15648298 | pseudogene | INBRED, LOCO | I:4443624, I:4443624 | 21, 21 | 4464670 | Yes | No |
| *Y105C5A.25* | *Y105C5A.25* | IV:15860221-15861442 | pseudogene | INBRED, LOCO | I:4446767, I:4446767 | 21, 21 | 4464670 | Yes | No |
| *Y59H11AR.6* | *Y59H11AR.6* | IV: 8597762-8598727 | pseudogene | INBRED, LOCO | I:4443624, I:4443624 | 21, 21 | 4464670 | Yes | No |
| *Y82E9BL.18.1* | *Y82E9BL.18* | III: 1328590-1331066 | protein-coding | INBRED, LOCO | I:4316016, I:4316016 | 20.5, 20.5 | 4464670 |  | Yes |
| *Y82E9BL.18.2* | *Y82E9BL.18* | III: 1326791-1331066 | protein-coding | INBRED, LOCO | I:4434616, I:4434616 | 21,21 | 4464670 |  | Yes |
| *ZK795.6* | *ZK795.6* | IV: 12555311-12556482 | pseudogene | INBRED, LOCO | I:4443624, I:4443624 | 21,21 | 4464670 |  | No |

**Extended Data Fig. 1.**
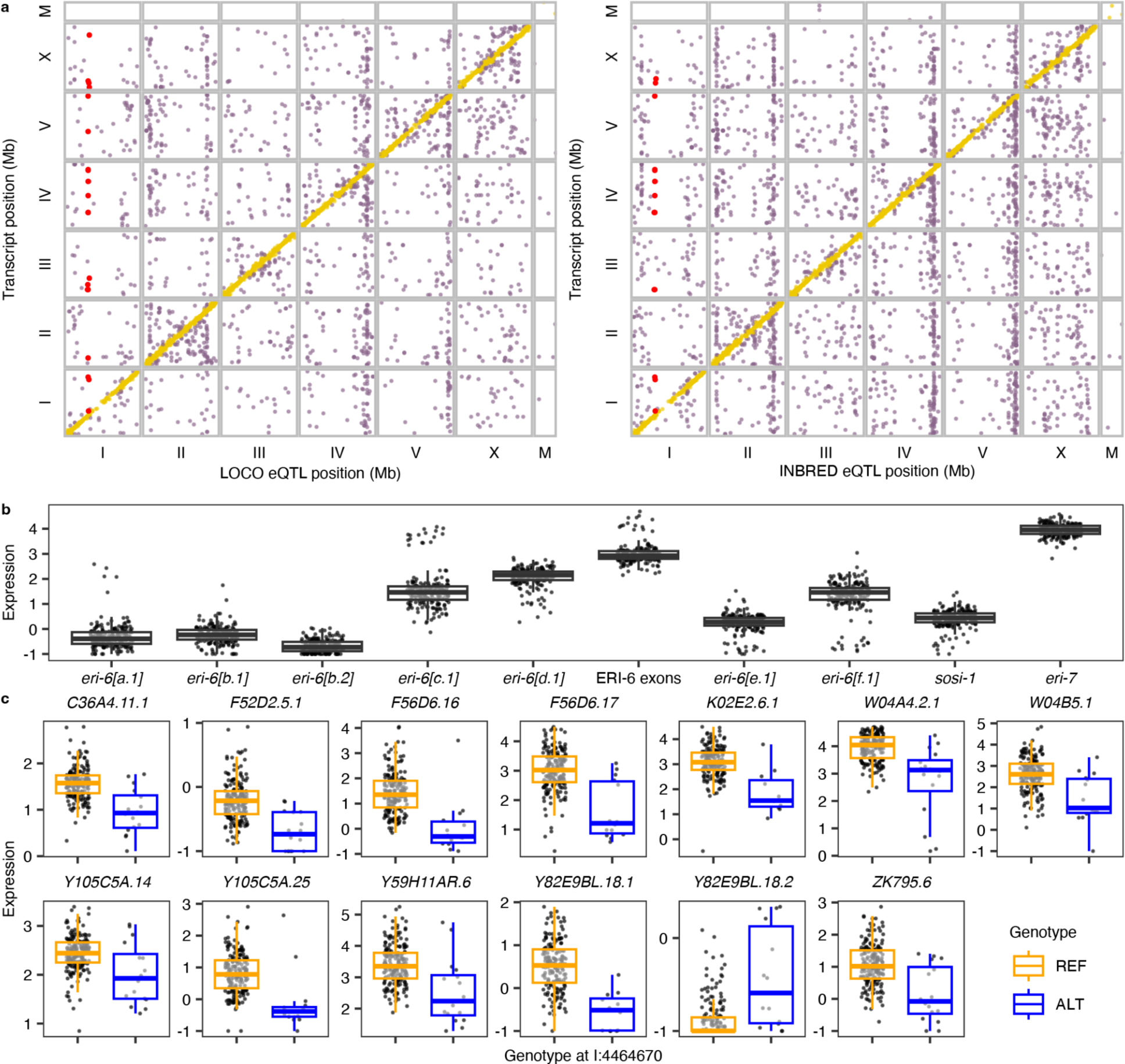
Expression QTL maps and expression variation of *eri-6* related transcripts. **a**, Expression QTL maps using LOCO and INBRED methods. Each point represents an eQTL with its position on the x-axis and the genomic position of the transcript on the y-axis. Local and distant eQTL are colored gold and purple, respectively. Red points represent distant eQTL associated with the *eri-6/7* locus. **b**, Tukey box plots showing expression (-*log*_2_(normalized TPM+0.5) variation of ten transcripts at the *eri-6/7* locus. **c,** Tukey box plots showing expression variation of 13 transcripts across the genome between strains with the reference (REF) or alternative (ALT) alleles at the SNV of 4,464,670 bp on chromosome I. **b,c,** Each point represents a strain. Box edges denote the 25th and 75th quantiles of the data; and whiskers represent 1.5× the interquartile range.

**Extended Data Fig. 2.**
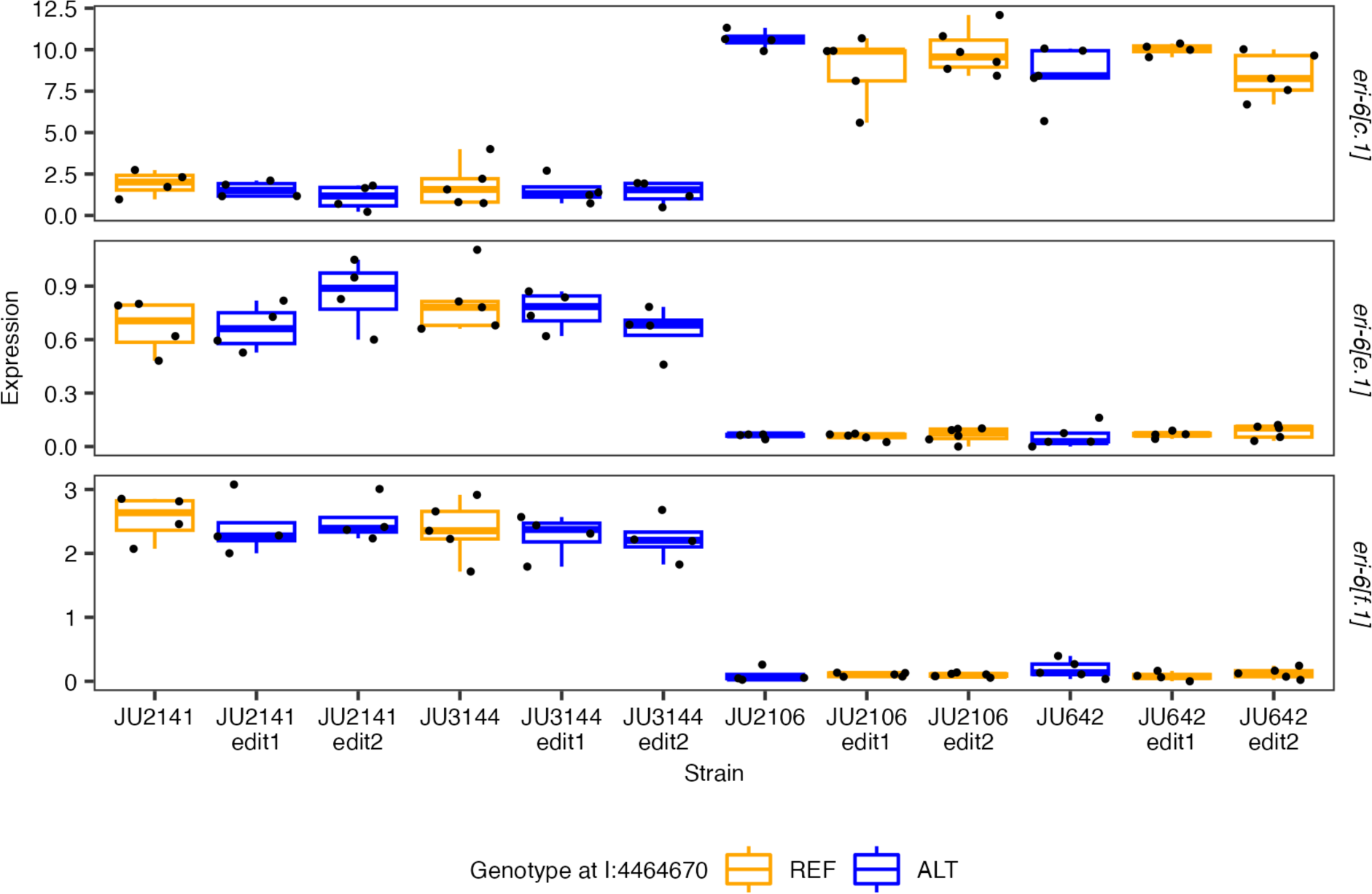
The SNV candidate cannot explain expression variation in *eri-6*. Expression variation of *eri-6[c.1,e.1,f.1]* among four wild *C. elegans* strains (JU2141, JU3144, JU2106, JU642) and their eight mutant strains at *eri-6[e]* (I: 4,464,670) using CRISPR-Cas9-mediated genome editing as previously described^1–3^. The guide RNAs crECA163 (GCTGTGCCACGATCGGAGTA) (Synthego, CA, USA) was used for the editing. The homologous recombination templates crECA162 (tgtcatttgatcccgctcggcattttcaacgatgacgaaaagtcttctaacatctcgaatTaccttactccgatcgtggcacagctcaatagcctcaaagagctgaaactgaaagtagccg) and crECA164 (tgtcatttgatcccgctcggcattttcaacgatgacgaaaagtcttctaacatctcgaatGaccttactccgatcgtggcacagctcaatagcctcaaagagctgaaactgaaagtagccg) (IDT, IL, USA) were used for wild strains with the reference (REF) and alternative (ALT) alleles at the target site, respectively. Genotypes of F2 progeny were detected with primers oECA1989 (GGTGGTGGCAGCGCATCTAGTC) and oECA1990 (GCTCCCCGAATGTAGCCACCGA) using PCR and Sanger sequencing. Edit1 and Edit2 are two independent edits in each of the four backgrounds. Each point represents a biological replicate. Transcriptomes in synchronized young adult stage animals of each replicate were measured by RNA sequencing and quantified as previously described^4^ (also see details in Methods).

**Extended Data Fig. 3.**
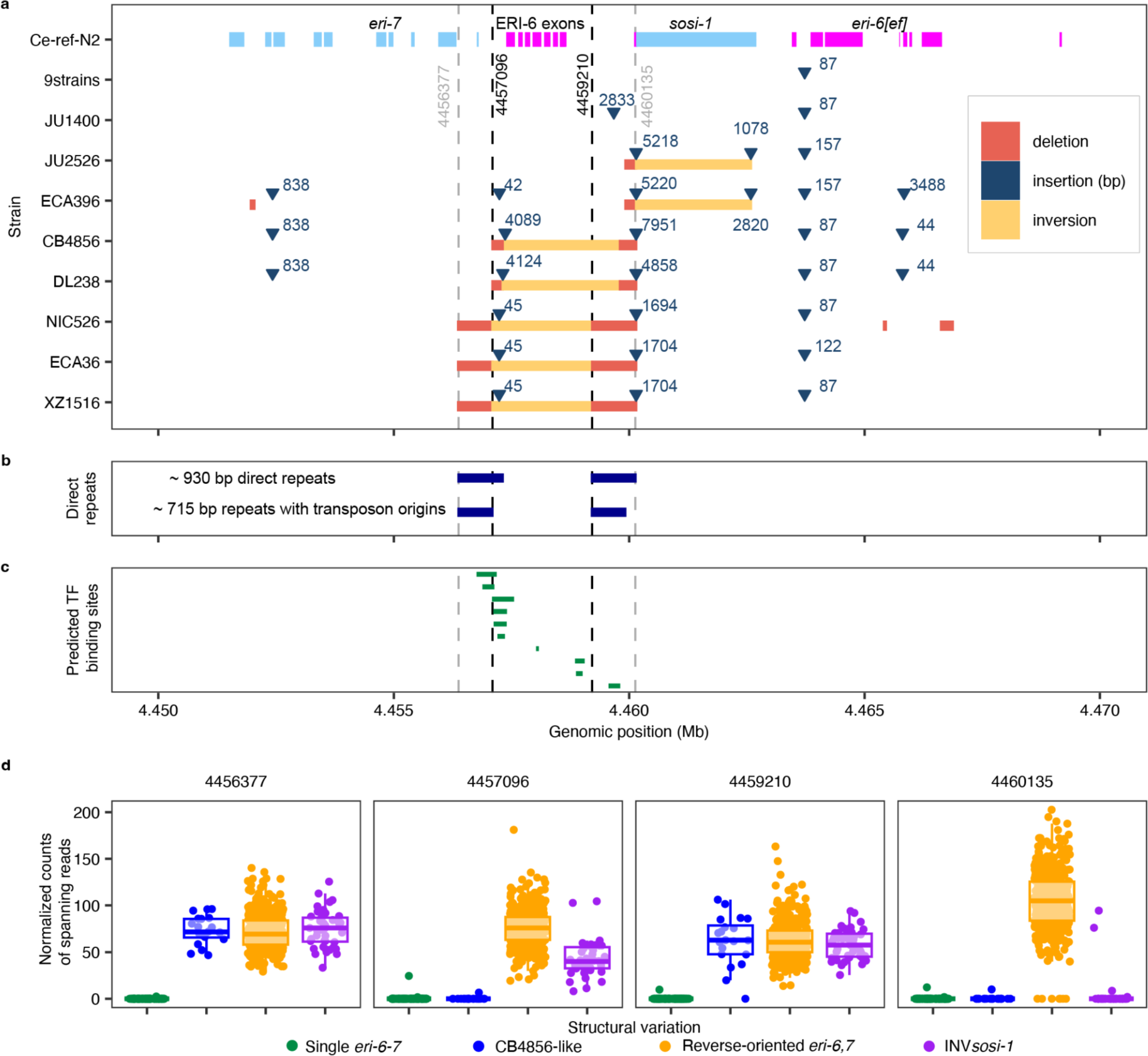
Structural variation in the *eri-6/7* region. **a**, Large deletions (≥38 bp), insertions (≥42 bp), and inversions of 17 wild strains with PacBio assemblies to the reference N2 genome in the *eri-6/7* region are represented by red rectangles, dark blue triangles, and yellow rectangles, respectively. Sizes of the insertions are indicated in bp. Exons of *eri-6/7* and *sosi-1* are plotted as rectangles on top and are colored magenta and light blue for plus and minus coding strands, respectively. Nine strains (DL226, EG4725, JU310, JU1395, JU2600, MY2147, MY2693, NIC2, QX1794) with highly identical sequences in this region were represented together. The only local structural difference in these nine strains as compared to N2 is a shared 87-bp insertion upstream of *eri-6[e]*. **b**, Ranges of the ∼930 bp direct repeats^5^ and the ∼715 bp parts with *Polinton* origins are indicated as blue rectangles, respectively (Supplementary Table 4). Dashed vertical gray and black lines indicate outside boundaries of direct repeats and break points of inversions (defined by comparison between the strains XZ1516 and the reference N2). **c**, Predicted transcription factor (TF) binding sites^6^ within the repeat regions are indicated as green rectangles. **d,** Number of reads spanning 20 bp surrounding each boundary/break-point position was counted and percent of this count normalized by the mean coverage per bp in the *eri-6/7* locus for each strain was plotted on the y-axis against the structural variation on the x-axis. High counts of reads spanning the inversion breakpoints indicate Watson ERI*-*6 exons and direct repeats as in the reference genome, except for the CB4856-like strains.

**Extended Data Fig. 4.**
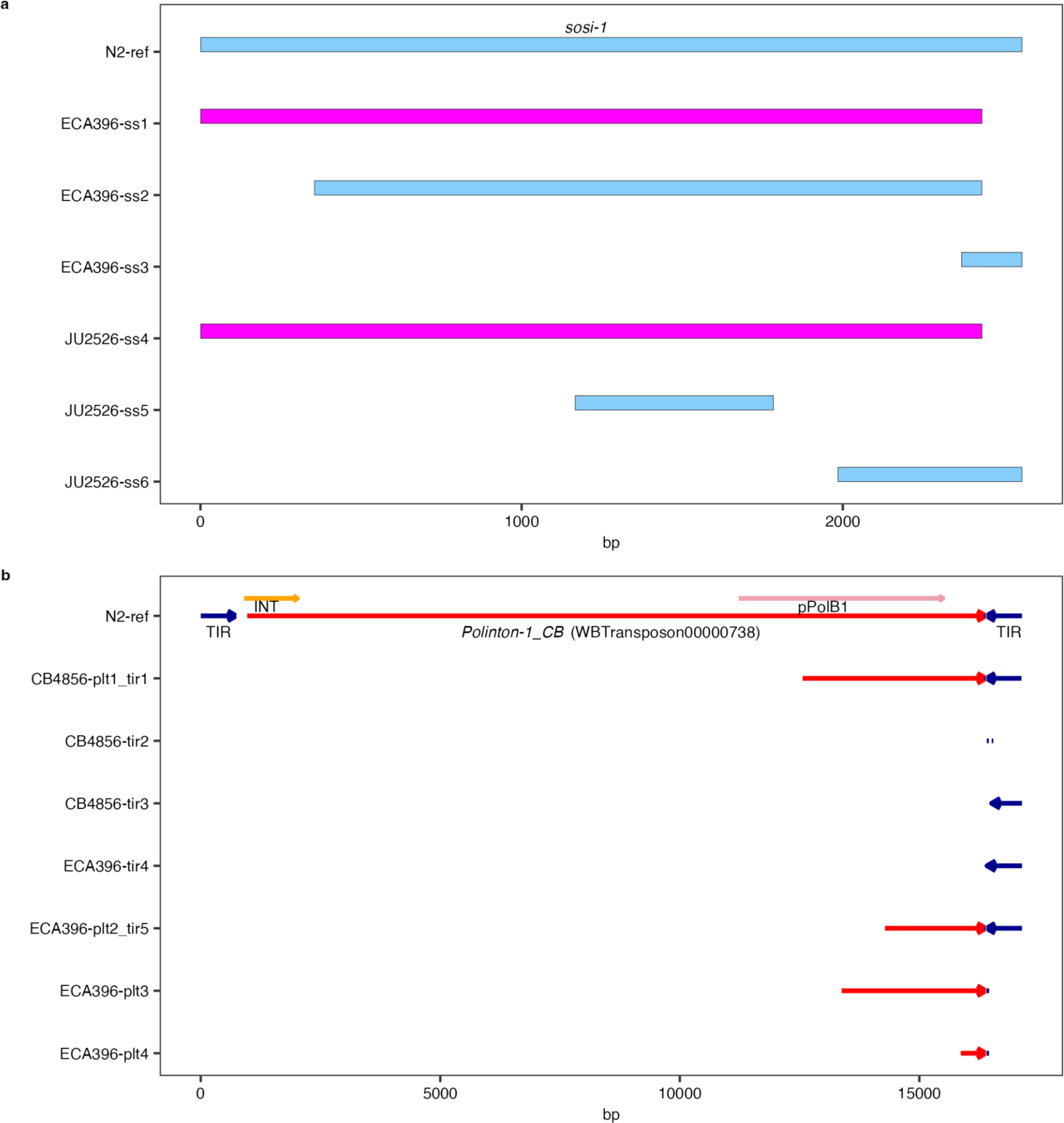
Alignment of *Polinton* and *sosi-1* fragments to the reference. Sequence alignments of fragments (“ss1-6”, “tir1-5”, and “plt1-4”) indicated in Fig. 2 to *sosi-1* (**a**) and the largest *Polinton* remnant (*Polinton-1_CB,* WBTransposon00000738, as a red arrow on top) (**b**) in the reference N2 genome are shown. Positions of the retroviral-like-element integrase (INT) and the protein-primed DNA polymerase B genes (pPolB1) are indicated as orange and pink arrows, respectively. Note that sequences of segments CB4856-tir2, tir3, and ECA396-tir4 can also be aligned to the terminal inverted repeat (TIR, blue arrows) on the left.

**Extended Data Fig. 5.**
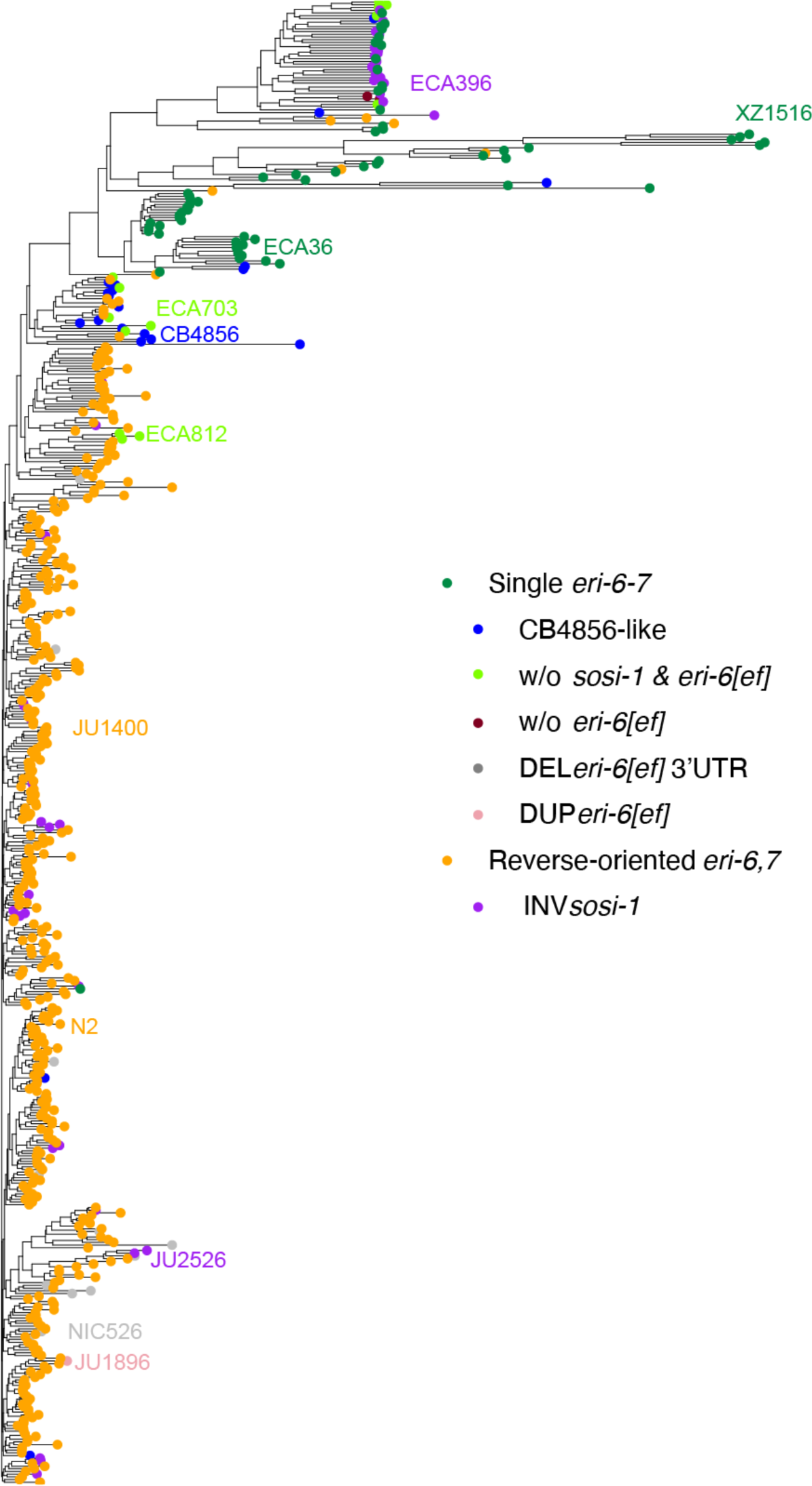
Genetic relatedness of 550 wild *C. elegans* strains. Genetic relatedness tree using 1,199,944 biallelic SNVs throughout the genome. Recombination occurs within the species so this tree only represents overall relatedness. Each point represents a strain and is colored by its structural variation in the *eri-6/7* region. The 19 strains clustered with ECA396 in Fig. 3 are also clustered with ECA396 on the same branch.

**Extended Data Fig. 6.**
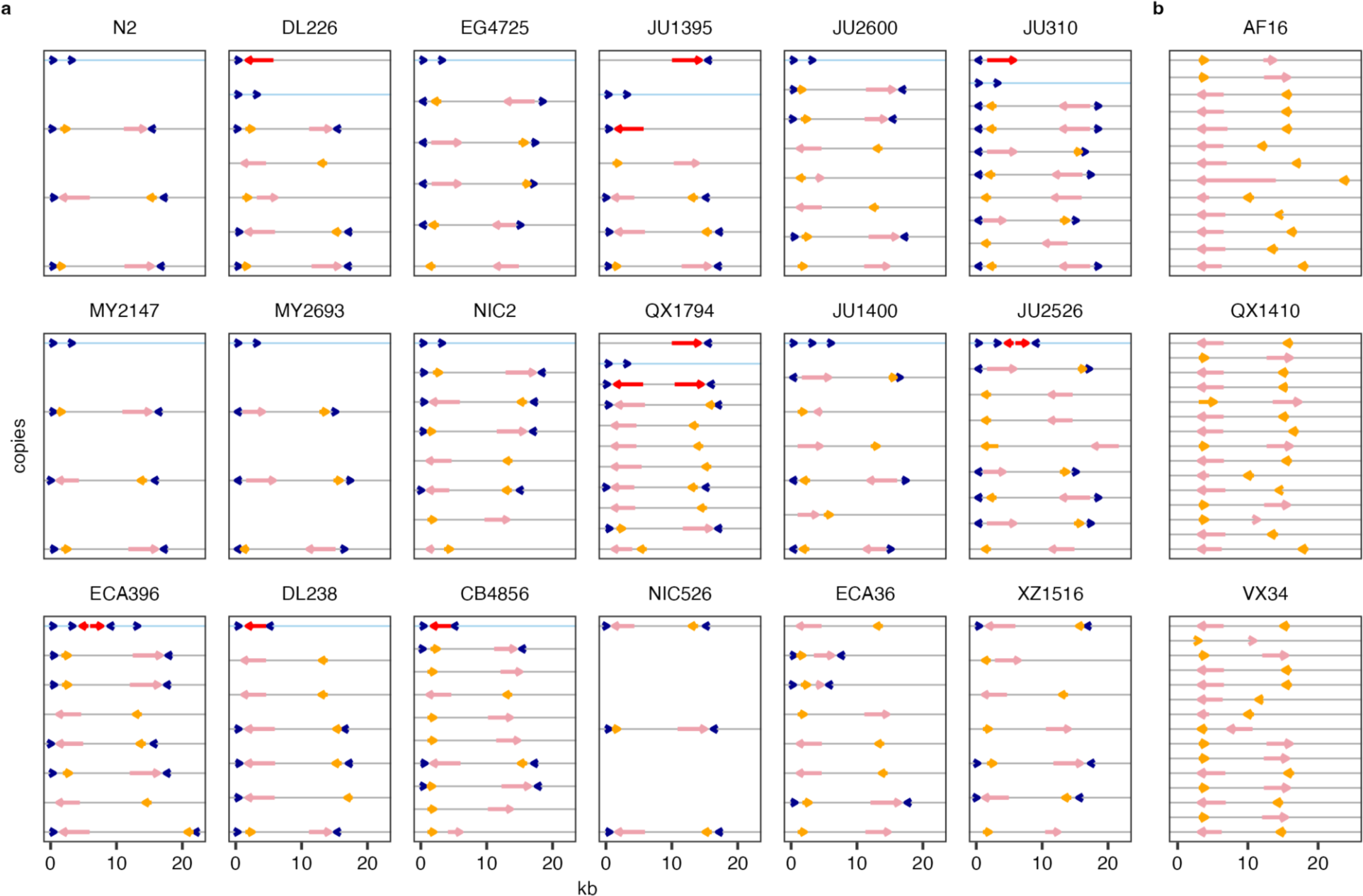
Presence of TIRs and *Polintons* in *C. elegans* and *C. briggsae* strains. *Polinton* insertions were identified in 18 *C. elegans* strains (**a**) and three *C. briggsae* strains (**b**) by requiring the presence of both pPolB1 (pink arrows) and INT (orange arrows) within 20 kb. Blue arrows represent TIRs. Red arrows represent pPolB that was found close to TIRs but without nearby INT. Direction of arrows represents orientations. Horizontal lines indicate different copies in the genomes, with the blue lines highlighting the copies in the *eri-6/7* locus. All the identified *Polinton* insertions and TIRs were plotted.

**Extended Data Fig. 7.**
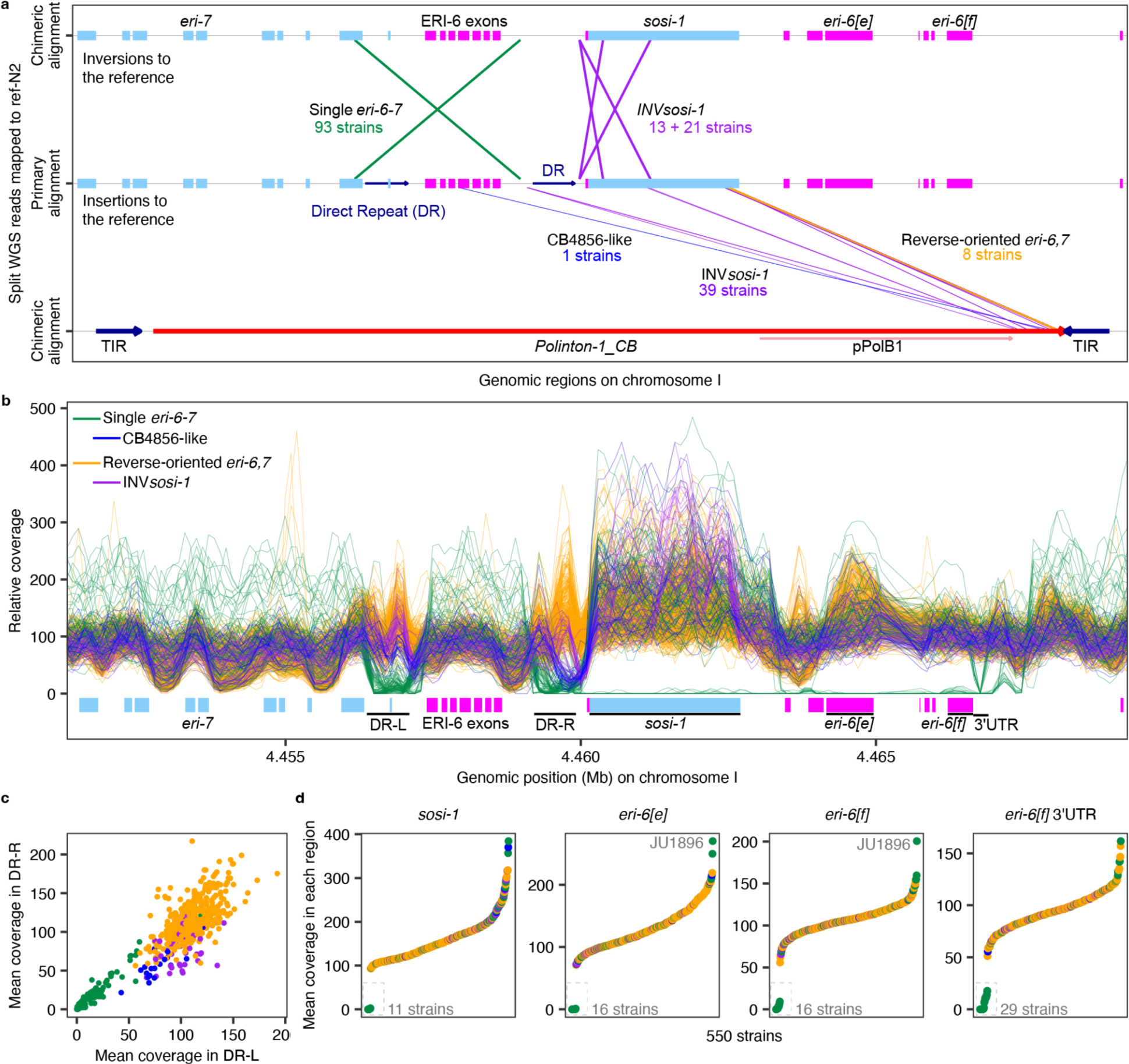
Inversions and other structural variants within the *eri-6/7* locus. **a**, Inversions and *Polinton-1_CB* insertions within the *eri-6/7* locus are represented by lines that connect positions of the primary and chimeric alignments of split reads (See Methods). **b**, Sliding windows on normalized coverages per bp with a 200-kb window size and a 100-bp step size in the *eri-6/7* locus of 550 strains. **c**,**d** Mean coverages in each strain within the two direct repeats (**c**) and four other regions (**d**) indicated in **b** were shown. Each line (**b**) / point (**c,d**) represents one strain and is colored by structural variants indicated in **b**. Extreme low coverages indicate deletions and extreme high coverages indicate duplications compared to the reference genome.

**Extended Data Fig. 8.**
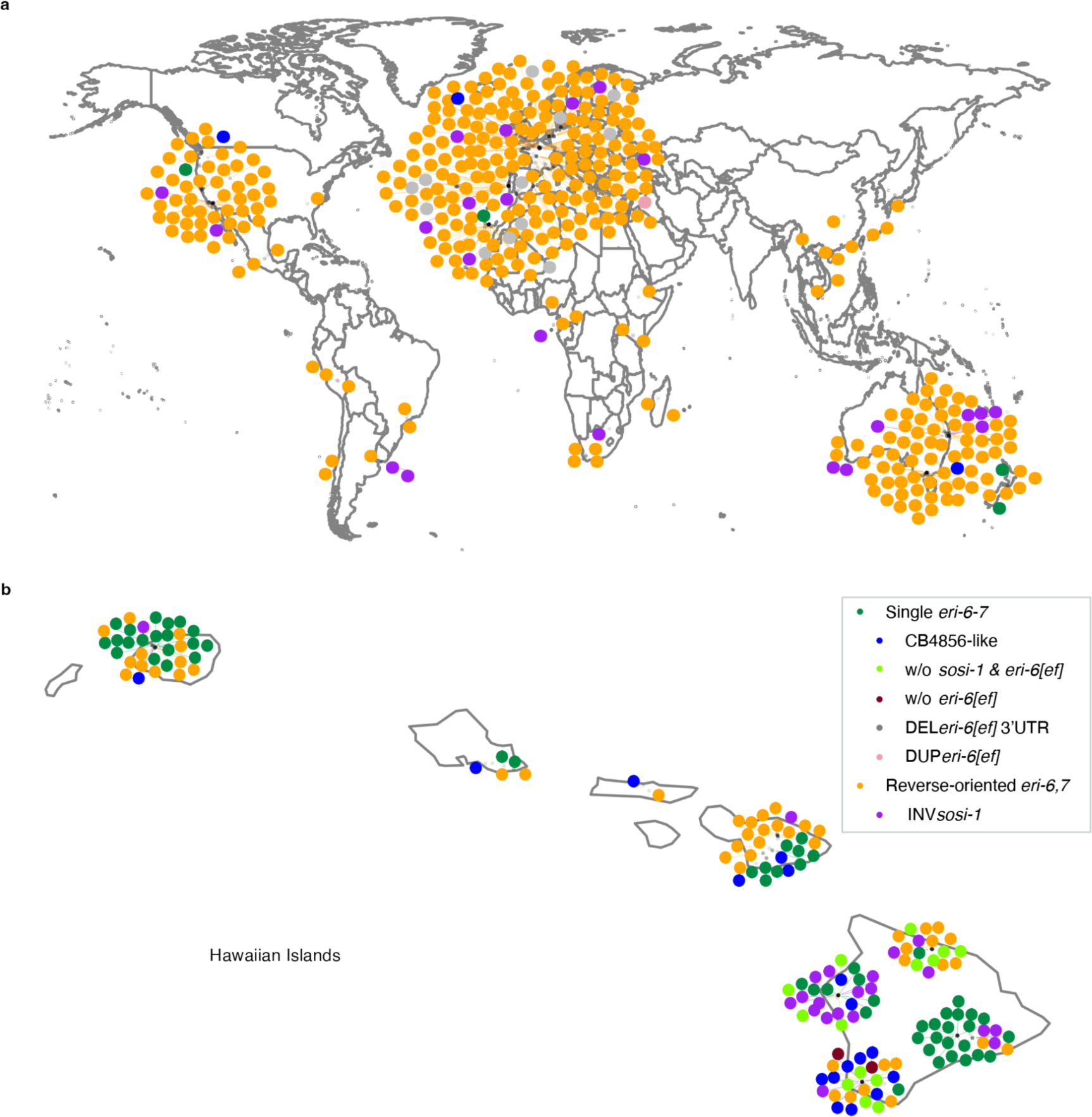
Geographical distribution of wild *C. elegans*. Geographical distribution of 550 strains worldwide (**a**) and detailed on the Hawaiian Islands (**b**). Each point represents a strain and is colored by its structural variation in the *eri-6/7* region.

